# Complementing Biosafety Levels With Societal Safety Levels

**DOI:** 10.64898/2026.10.28.762599

**Authors:** Kevin M. Esvelt, Dana Gretton, Lennart Justen

## Abstract

Biosafety level (BSL) classifications of biological agents have succeeded at minimizing occupational risk inside laboratories, but largely neglect the potential for onward transmission. As a result, many non-transmissible pathogens that could kill at most one individual per person exposed require greater containment than 1918 influenza, a virus that killed 50-100 million and might cause a new pandemic today. Laboratory workers are consenting professionals; the public is not. We recommend complementing BSLs with explicit thresholds for expected societal harm. Here we formalize Societal Safety Levels (SSLs) based on expected public fatalities per year, which we quantify from (i) estimated probability of laboratory release, (ii) likelihood of societal transmission, and (iii) consequence. SSLs would apply specifically to laboratory research with replication-competent potential pandemic pathogens, and would not restrict field surveillance, sequencing, or countermeasure development using non-replicating systems or pandemic-incapable agents. We find that applying the 0.1 expected deaths per facility-year standard common across multiple major-hazard industries to potential pandemic pathogen research would require marked safety improvements. Complementing BSLs with SSLs would establish enforceable accident- safety thresholds for pandemic-capable pathogens, bringing high-risk biological research into alignment with the quantitative safety standards that help other hazardous industries operate with public acceptance.

## Introduction

According to Biosafety in Microbiological and Biomedical Laboratories (BMBL, 6th Edition), "The fundamental objective of any biosafety program is… to minimize the exposure of laboratory personnel, the public, and the environment to biological agents"[1]. Ongoing disputes over the safety of enhanced- transmission research and the proximal origin of SARS-CoV-2 have intensified scrutiny of whether current classification practices adequately fulfill this public protection mission, damaging trust in science[2–9]. Regardless of whether inadequately regulated wildlife markets or unsafe research activities were responsible for COVID-19, it is clear that regaining public trust will require the scientific community to transparently monitor and mitigate risks to the public using the same quantitative rigor that we apply to our own research.

Effective public protection requires biosafety ratings to quantify the expected harms from working with each biological agent and mandate safeguards proportionate to increasingly severe risks. Therefore, experiments involving transmissible biological agents should require greater laboratory safeguards than work with non-transmissible pathogens that can only harm researchers who have been directly exposed in the laboratory.

The available evidence suggests that current BSL classifications, while effective at protecting laboratory workers, fail to adequately account for onward transmission. The 1977 H1N1 epidemic, widely accepted to have resulted from human error because the genome of the virus was virtually identical to a 1950s strain, killed an estimated 700,000 people, conclusively demonstrating that laboratory strains can spread globally[10–12]. Yet experiments involving the 1918 H1N1 influenza virus, which killed between 50 and 100 million people and is thought to present a nontrivial risk of a new pandemic today, require fewer precautions than many studies involving non-transmissible agents that cannot possibly harm anyone outside of the laboratory[1]. If systematic assessment were to show that this example represents a general pattern – greater containment for agents that cannot spread beyond a single individual than for agents capable of infecting billions – it would constitute strong evidence that the current biosafety classification system employed by the BMBL (Risk Group assignments and related BSL requirements) fails to adhere to its stated intent of minimizing public exposure.

Previous analyses have performed limited quantitative risk assessments of research with enhanced potential pandemic pathogens and explored modeling approaches to estimate accident probabilities[4,5]. However, the first systematic empirical data on laboratory incident rates specifically involving community exposure potential has only recently become available through Freedom of Information Act releases of Federal Select Agent Program and NIH records[13]. These datasets have limitations: incident reports are heterogeneous, denominators are imprecise, and rates may reflect both over-counting (inclusion of events unlikely to cause community transmission) and under-counting (unreported incidents). Here we use the dataset with the lowest entity-level community release estimate to translate observed rates into expected harms, perform sensitivity analyses, and propose Societal Safety Levels (SSLs), a tiered regulatory framework aligned with established risk assessment practices.

SSLs would adopt the regulatory logic used in chemical, hydropower, aviation, and other major-hazard industries by establishing fixed, quantitative risk thresholds that activities must not exceed. These industries limit the acceptable individual risk of death (IR) for each person in the exposed community. Aggregating this risk over the impacted population enables direct comparisons of expected deaths per facility-year. The SSL framework would extend this logic to transmissible agents with the potential to cause a new outbreak (and consequently would *not* apply to currently circulating or recent endemic pathogens such as seasonal influenza). By establishing similar thresholds and requiring enhanced precautionary protocols to keep risks tolerable, SSLs would complement existing worker-focused BSLs to better protect local and global communities.

The July 2026 U.S. Government Policy for Stopping High-Risk Life Sciences Research prohibits dangerous gain-of-function research and requires risk-benefit review of research with the potential to produce such outcomes[14]. However, the policy doesn’t establish quantitative thresholds for accidental harm imposed on the public. SSLs are intended to fill this gap. Throughout this analysis, ’societal risk’ refers to expected public harm from accidental release and onward transmission.

## Results

### Current biosafety classification protects workers but neglects community risk

Using site counts and personnel numbers consistent with current practice, we estimated today’s frequent BSL-4 workers at roughly 500-1,500 worldwide, and total authorized personnel at 3,000-10,000 (Methods). Combined with the fact that only a single person has died from a laboratory-acquired infection involving a BSL-4 pathogen since 2000[15], the fatal infection risk from strict BSL-4 procedural work is on the order of ∼0.3–6 per 100,000 worker-years. While the available data are limited, this is comparable to or lower than most industrial occupations, indicating that BSL-4 precautions are effective at protecting workers[16].

However, these same biosafety classifications may not adequately mitigate community risk stemming from outbreaks of transmissible pathogens, which can cause harm to the general population in the event of an accidental release. To determine how well current biosafety classifications reflect outbreak risk, we compiled a table of the largest natural outbreaks of each BSL-4 pathogen and compared them to the five most damaging natural outbreaks of any pathogen in recent history (Supplementary Table S1).

Excluding variola (continuously endemic until eradication), the largest outbreak from an agent that would typically require BSL-4 containment caused 11,325 deaths: the 2014-16 West Africa Ebola epidemic (Fig. 1a)[17]. Summing the largest outbreaks of every remaining BSL-4 agent yields fewer than 1,000 additional deaths, mostly in settings with fragile health-care systems(Supplementary Table S1). BSL-4 laboratories are heavily concentrated in high-income nations with robust medical surveillance and infection control[18,19]; most epidemiologists are deeply skeptical that Ebola or other highly lethal non-respiratory pathogens could spread widely in such settings[20–22]. Therefore, laboratory accidents involving non-variola, wild-type BSL-4 agents appear to pose limited risk to local communities or the global public.

**Figure 1.**
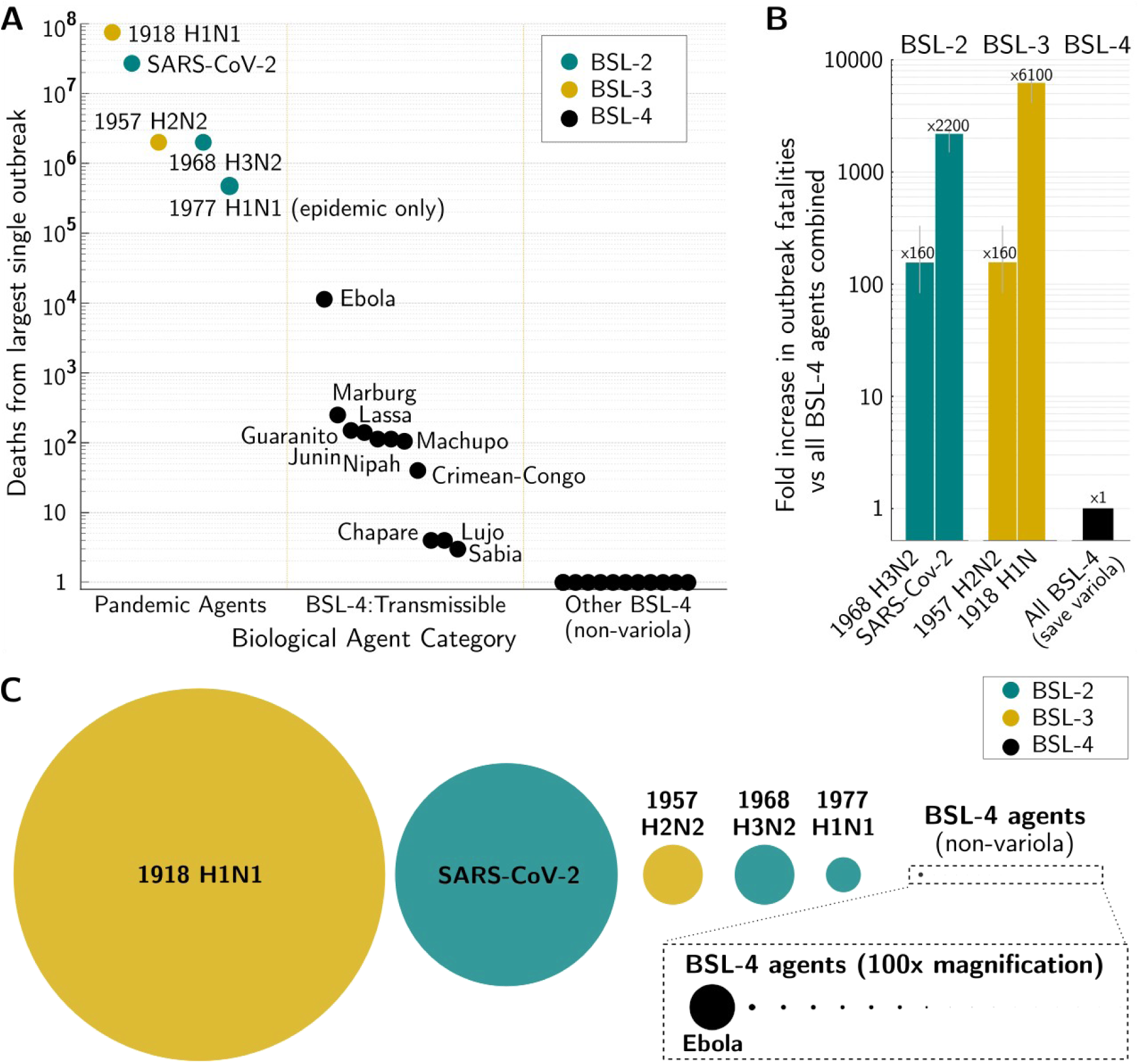
**| Biosafety containment requirements neglect community risks from outbreaks of pandemic agents**. **a)** Death tolls from the largest recorded outbreaks of BSL-4 agents (excluding smallpox, which was globally endemic) are far smaller than outbreaks of pandemic agents, yet non-variola potential pandemic pathogens from any source require only BSL-2 or BSL-3 containment. **b)** Historical pandemics have killed orders of magnitude more people than all non-variola BSL-4 outbreaks combined, demonstrating that current biosafety classifications effectively protect workers but inadequately account for risks to the general public from potential pandemic agents. c) Bubble plot of outbreak death tolls to clarify the scale difference. Even the anthropogenic 1977 H1N1 epidemic, not usually considered a pandemic, killed an estimated 700,000 people: more than 50 times the sum of the largest outbreaks from each BSL-4 agent.

By contrast, more transmissible but less lethal agents have caused far greater harm. Research with these pathogens and agents suspected of similar capabilities requires many fewer safeguards. 1918 H1N1 influenza (BSL-3+) killed 50-100 million. SARS-CoV-2 (BSL-2) killed 27 million. 1957 H2N2 influenza (BSL-3) and 1968 H3N2 influenza (BSL-2) each killed approximately 2 million[23]. The 1977 H1N1 influenza epidemic (BSL-2) killed an estimated 700,000 after being out of circulation for just 25 years [10,11].

The death tolls of these historical respiratory outbreaks exceeded the casualty counts of non-variola BSL-4 agents by two to five orders of magnitude, indicating that transmissible but less lethal agents pose a far greater risk to the general public (Fig. 1a-b).

Even the estimated death toll of the relatively mild 1977 H1N1 epidemic[24,25] suggests that the originating accident plausibly killed twenty times as many people as the combined toll of every non-variola BSL-4 agent across all of modern history. Yet these historical agents, as well as zoonotic potential pandemic pathogens and products of enhanced-transmission research, require only BSL-2 or BSL-3 containment. While some research on potential pandemic pathogens is conducted at ’BSL-3+,’ this designation describes a heterogeneous set of ad hoc practices less stringent than BSL-4. The gap suggests that current biosafety ratings fail to mitigate the most severe risks to the general public.

One might argue that historical pandemic pathogens pose negligible risks today due to population immunity gained from exposure to their modern descendants. This is almost certainly true for SARS-CoV-2, given how recently it spilled over and infected virtually every human. However, as immunity to historical viruses wanes with time, these demonstrably pandemic-capable agents increasingly risk new outbreaks; declining immunity explains why the 1977 H1N1 epidemic primarily infected younger people who had never been exposed[26]. This risk warrants greater caution than is reflected by current biosafety levels. For example, while current immunity to the 2009 H1N1 virus and its descendants provides some protection against 1918 influenza[27,28], the potential consequences of that virus spreading today are severe enough that any nontrivial probability demands quantitative risk assessment rather than dismissal.

The inadequacy of the current biosafety classification system with respect to potential pandemic pathogens is even clearer when considering zoonotic research. Two coronaviruses have proven capable of spreading widely in humans, out of just a few hundred bat coronaviruses in nature thought capable of infecting human cells[29,30]. Laboratory studies of these newly discovered zoonotic agents are routinely conducted at BSL-2 or BSL-3 depending on the nation – never BSL-4[29]. Yet SARS-CoV-2 killed more than 2,000 times as many people as the combined worst outbreaks of every BSL-4 agent (excluding previously endemic variola). These numbers indicate that any single bat CoV deemed worthy of studying because it is thought to pose a greater risk represents a threat comparable to if not exceeding the entire non-variola BSL-4 pathogen collection, yet experiments require less reliable containment.

Having demonstrated that current BSL ratings appear to adequately protect laboratory workers while largely neglecting the possibility of onward transmission through communities, we next considered how best to formulate a complementary system to manage risks to the public while preserving the concrete benefits of medical research.

### Societal Safety Levels quantify expected public harm

We propose to safeguard society from transmissible biological agents studied in laboratories by establishing Societal Safety Levels (SSLs) based on quantitative thresholds for expected public harm. Research with high-SSL pathogens would require escalating oversight to keep total risk to communities below a broadly acceptable level. Only high-consequence agents would be impacted: the framework would not apply to prions, toxins, attenuated lab strains (H1N1 PR-8), endemic pathogens circulating in humans (SARS-CoV-2), or zoonotic agents causing frequent human infections without sustained transmission (wild-type H5N1).

**SSL-1 (Minimal risk)**: No additional safeguards required; current BSL precautions suffice.

**SSL-2 (Low risk)**: Add mandatory public incident reporting and annual safety reviews.

**SSL-3 (Moderate risk)**: As SSL-2, plus personnel screening and a concrete benefits review.

**SSL-4 (High risk)**: Requires an independent board evaluating a quantitative risk-benefit analysis to conclude that concrete expected benefits will greatly exceed harms.

**SSL-5 (Extreme risk)**: Analogous to smallpox: requires specific authorization by international resolution (e.g., World Health Assembly acting under U.N. Security Council mandate)

SSL-1 through SSL-4 parallel the BSL numbering system, but are complementary: where BSLs demonstrably protect consenting laboratory professionals, SSLs protect the general public. When working with a particular agent, the researcher would conform to both BSL and SSL requirements. The addition of SSLs would mandate additional safeguards for certain high-risk transmissible pathogens, especially those with pandemic potential, without impacting the vast majority of microbiology research.

We structured these categories using order-of-magnitude increases in expected harm, with each SSL tier representing roughly a tenfold increase from the previous level (Fig. 2). This enables systematic risk assessment across research portfolios and provides clear decision criteria. Crucially, SSLs update as epidemiological conditions change. When an agent achieves widespread human circulation, the marginal contribution of laboratory releases to total disease burden approaches zero. SARS-CoV-2 transitioned from high-SSL (pandemic threat) to SSL-1 (endemic) within weeks of global spread. Conversely, waning population immunity may increase SSL over time, as occurred for 1977 H1N1 after 25 years out of circulation. This bidirectionality is intentional: SSLs track current societal risk from accidental releases, not static hazard classifications.

**Figure 2.**
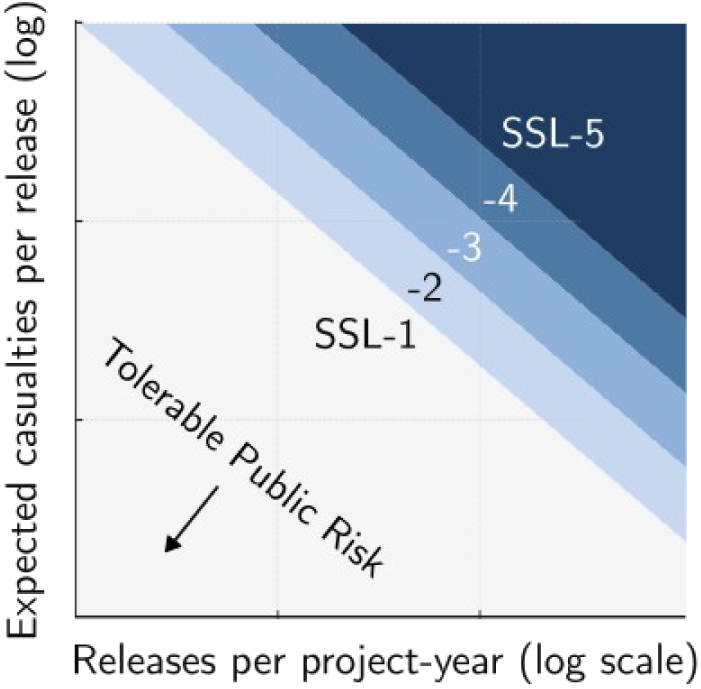
| Societal Safety Levels would keep expected harm below a tolerable threshold. To keep expected harm below a threshold, the likelihood of a hypothetical laboratory-originated pandemic must decrease in proportion to increasing pathogen severity. Therefore, Societal Safety Level increases with expected casualties, chance of pandemic given a community release, and frequency of community releases. More lethal (right panel) or contagious (higher Y axis value) viruses require superior containment (lower X axis value) to meet a given SSL. SSL-5 would represent presumptively intolerable risk, the level of which we subsequently determine by considering standards across other industries.

To implement this framework, we developed methods to quantify expected harm to society from laboratory accidents involving transmissible pathogens.

### Quantifying expected public harms from community releases of transmissible agents

In order to assign SSLs consistent with risks tolerated across other industries, we developed a framework to quantify expected harms to the general public resulting from laboratory research involving transmissible pathogens. This framework incorporates accident rates that result in community exposure, epidemiological parameters of the pathogen, and response effectiveness that collectively determine likely outbreak trajectories.

A laboratory release occurs when an individual harboring an undetected or unreported laboratory-acquired infection with a transmissible agent interacts with the community.

● **p_init_** = probability of an undetected or unreported laboratory-acquired infection leading to community exposure, events per year (a researcher may seed an outbreak)

#### Laboratory accident rates leading to community exposure

We estimate accident probabilities using two FOIA-obtained U.S. datasets tracking undetected or unreported laboratory-acquired infections that presumably resulted in community exposure[13]:

● **Baseline estimate (FSAP data, low-end)**: 0.25% chance per year
● **Higher estimate (NIH BSL3/4 data, low-end)**: 1.6% chance per year

These datasets have noteworthy limitations. Both refer to entities, which may have multiple labs working with one eligible agent, multiple agents, or conduct less than one full year of work per registered agent. Incident reports are heterogeneous, and denominators for total experiment-hours and risk categories across institutions remain imprecise. Reporting requirements formally extend to near-misses and equipment failures, but compliance likely varies across institutions, and asymptomatic or mild laboratory-acquired infections may go unrecognized regardless of reporting culture. The degree to which current data over- or undercounts true community exposure risk remains uncertain. We recommend updating Form 3 to request structured incident details under no-fault reporting, modeled on the Federal Aviation Administration’s Aviation Safety Reporting System, to clarify this uncertainty.

Throughout this analysis, we use the lower FSAP-derived estimate (0.25%) as our baseline. As more laboratories conduct similar research, the probability that at least one accident occurs increases proportionately[18].

#### Outbreak trajectories of transmissible pathogens

Each community release produces one of three outcomes (Figure 3):

The transmissible agent fails to infect anyone else (e.g. SARS, Singapore 2003)
A local outbreak that soon burns out and goes extinct (e.g. SARS, Beijing 2004)
An epidemic or pandemic that spreads worldwide (e.g. 1977 H1N1)

**Figure 3.**
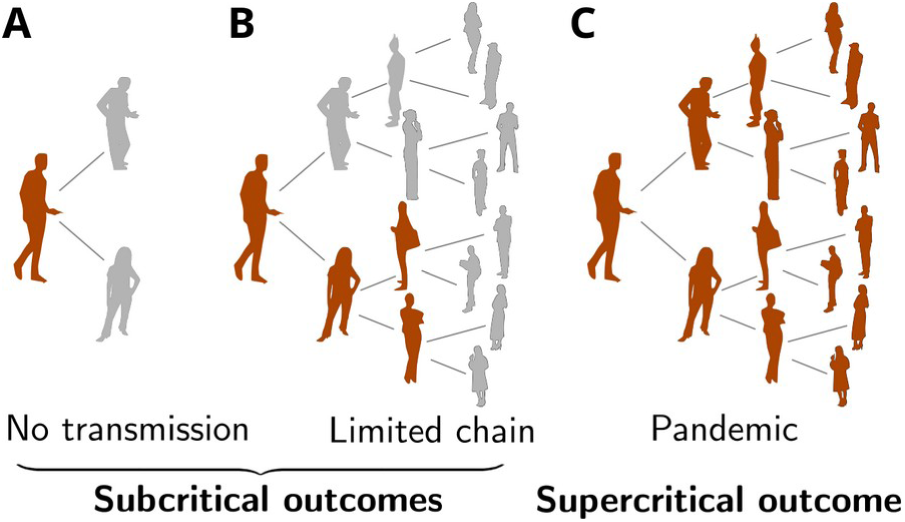
| Potential outcomes from community releases of transmissible agents. (A) No transmission, (B) limited chain extinction, or (C) pandemic. Subcritical agents (*R_eff_* < 1) produce only A and B; supercritical agents (*R_eff_* > 1) can produce any outcome, but expected harm is dominated by outcome C.

#### Exempt agents

Two categories of agents pose negligible risk of limited-chain and especially pandemic outbreaks. All recent and currently circulating strains of endemic human pathogens are blocked by existing population immunity. Because the circulating wild-type strain is vastly more abundant than any potential release, laboratory accidents involving recent or current strains (e.g. seasonal influenza, SARS-CoV-2) would represent negligible contributions to the total burden of disease. Similarly, laboratory-adapted strains with demonstrated attenuation (e.g., Influenza A/PR/8/34) pose minimal risk of sustained transmission.

Therefore, both recent endemic strains and proven attenuated laboratory strains are exempt from SSL classification.

#### Subcritical agents (R_eff_ < 1)

A subcritical agent cannot sustain transmission indefinitely in the relevant population, so community releases are limited to outcomes A and B: local outbreaks that go extinct. We calculate expected deaths per year of research per laboratory using:

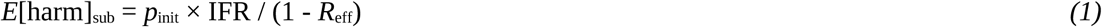

where IFR is the infection fatality rate and the term 1/(1 - *R*_eff_) represents the expected total number of infections per release event.

#### Supercritical agents (R_eff_ > 1)

Supercritical agents can produce all three outcomes. While outbreak extinction (outcomes A-B) remains possible, expected harm is negligible compared to epidemic or pandemic spread (outcome C). We therefore calculate expected deaths per year of research per laboratory as:

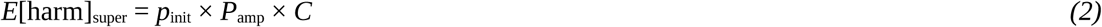

where:

● ***P*_amp_** = probability that a community exposure event amplifies into a large outbreak rather than dying out, calculated using established epidemiological methods[30]
● ***C*** = number of deaths if a large outbreak occurs

For agents with symptomatic presentation and high case fatality (variola, SARS-CoV-1), P_amp_ is substantially reduced by early detection and behavioral response in settings with robust surveillance. However, agents with mild or influenza-like symptoms pose substantially higher risk: unreported or undetected laboratory-acquired infections can be easily mistaken for seasonal respiratory illness, enabling multiple transmission generations before recognition. P_amp_ is correspondingly highest for agents with extensive asymptomatic transmission (SARS-CoV-2), as this trait renders containment much more challenging (Methods).

We estimate deaths using historical precedents scaled to the population that would be impacted by the release today. For novel pathogens that would establish as new endemic threats, we assume a large fraction of humanity would be infected: an equivalent of 1957 H2N2 would likely kill two to eight million, another SARS-CoV-2 approximately 27 million, and a new 1918 influenza would be limited to between 20 and 80 million (less than 1% IFR vs 2.5% historically) thanks to lingering immunity and modern therapeutic options[23].

Determining whether an agent is subcritical or supercritical requires assessing its transmission characteristics. As noted above, current or recently circulating strains of human pathogens such as seasonal influenza and SARS-CoV-2 pose negligible societal risk because they are already spreading through the population; for simplicity, we treat them as effectively subcritical. For well-studied pathogens with extensive human transmission data, this classification is straightforward based on observed R₀ values and transmission patterns. For agents with limited data, such as newly isolated zoonotic viruses or laboratory-generated variants, we use viral family transmission patterns and spillover evidence to assess pandemic potential (Methods). When insufficient evidence exists to classify an agent from a high-risk family as subcritical (fewer than 60 spillovers), but transmission parameters remain unknown, we define *π* _super_ as the probability the agent is capable of supercritical spread if it does not stochastically die out (*R*_eff_ > 1). We adopt 60 spillovers because a pathogen with P_amp_ = 0.22 has a probability of observing no sustained outbreaks of (1 – 0.22)^60^ = 3.4 × 10^-7^. Moreover, most spillovers are presumably undetected, and agents with near-threshold transmissibility can likely be controlled with behavioral change. Expected harm becomes:

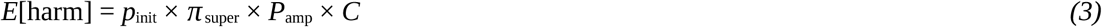

We conservatively estimate transmission parameters using the most transmissible well-characterized member of the same viral family when calculating *P*_amp_. Again, *π* _super_ reflects the probability that the agent would actually achieve sustained transmission. For agents lacking family-level precedents, such as primate arteriviruses[31], *P*_amp_ and *π* _super_ must be estimated from any limited spillover data and biological plausibility.

#### Contextual factors determining SSL assignment

Quantitative risk assessment provides a crucial foundation, but identical agents may warrant different SSL assignments depending on the experimental context. Relevant factors include procedure, local endemicity, facility infrastructure, and personnel experience. For zoonotic pathogens like H5N1 avian influenza, where natural human infections are frequent (SSL-1), laboratory mutagenesis or passage selecting for human adaptation may generate variants with pandemic potential exceeding wild-type. Such experiments warrant SSL assessment based on properties of plausible laboratory outputs, not naturally circulating strains. These contextual adjustments should be documented and justified in institutional review, not applied silently.

Geographic context similarly affects both baseline exposure and outbreak potential. In regions where an agent circulates naturally, laboratory work contributes negligibly to total community exposure; Ebola research in equatorial Africa adds little to baseline spillover risk. In high-income non-endemic regions, laboratories represent the primary exposure source, but robust surveillance limits outbreak potential. The highest-risk scenario may be laboratories in densely populated regions with limited healthcare infrastructure and no natural circulation - where the lab constitutes the sole exposure source and local R_eff_ may exceed estimates derived from high-income settings. SSL assignments should consequently be adjusted to account for facility location, rather than assume uniform global conditions. The framework provides a quantitative foundation for risk-informed decision-making, not a replacement for institutional judgment.

### Defining SSL thresholds based on established societal risk tolerances

Having established methods for calculating expected harms from releases, we next addressed the critical question of where to set the numerical boundaries between SSL categories in order to permit vital research while remaining worthy of public trust. We consequently looked to other industries for practical examples of risk management thresholds.

Regulators across diverse industries have converged on an individual risk (IR) of 10⁻⁶ deaths per person per year as the broadly acceptable limit for non-consenting members of the public exposed to major- hazard facilities[32–34]. This standard appears in dam safety guidelines, chemical facility regulations, and nuclear safety goals. The populations considered exposed to such facilities - determined by facility siting, impact radius, and urban geography - typically number around 100,000 people[35–38].

Aggregating this individual risk over the exposed population yields approximately 0.1 expected deaths per facility-year as the implicit threshold that major-hazard industries must not exceed.

Multiple established industries demonstrate this apparent convergence. In a representative example of dam safety, the Australian National Committee on Large Dams sets 10⁻⁶ individual risk as the broadly acceptable threshold[34]; applied to approximately 100,000 people downstream from major dam failure, this yields 0.1 expected deaths per dam-year[35]. Large chemical facilities and pipelines follow similar standards. For example, the UK Health and Safety Executive establishes 10⁻⁶ individual risk as broadly acceptable for the approximately hundred thousand residents within the impact radius of major petrochemical complexes such as Grangemouth, again yielding 0.1 expected deaths per facility-year[32]. Civilian nuclear power plants operate under even more stringent regulations, with U.S. Nuclear Regulatory Commission safety goals limiting individual risk of prompt fatality to 10⁻⁶ for populations of 10,000 or more within exclusion zones and severe accident impact areas, adjusted for background fatality risks[39].

Aviation provides complementary evidence based on observed performance rather than prescribed standards. In 2022, commercial aviation recorded 158 fatalities across 32.2 million flights, or 4.9×10⁻⁶ deaths per flight[40]. A regional airport handling 20,000 flights annually – comparable to Burlington, Bozeman, or Key West – would therefore experience approximately 0.1 expected deaths per airport-year.

The observed threshold of 0.1 deaths per facility-year is validated by examining facilities that exceed it. The failure of Warragamba Dam in Australia would place approximately 134,000 people on the downstream floodplain at risk, and accordingly has been the subject of decades of public safety debate and multi-billion dollar remediation proposals[35]. Paradise Dam was found to pose "unacceptable risk of dam failure" to downstream residents, a discovery that led to an independent Commission of Inquiry, an emergency reduction of the dam’s water level, and a decision to replace the dam entirely[41]. That facilities substantially exceeding 0.1 expected deaths per year are treated as public safety crises requiring regulatory intervention, public inquiries, and costly remediation supports the argument that this represents an upper limit for acceptable risk from beneficial but hazardous activities.

Research laboratories working with transmissible agents, like these other facilities, create benefits for society while imposing involuntary risks on surrounding populations. We therefore adopt this threshold as the boundary for SSL-5: research projects involving live transmissible pathogens capable of replicating in unmodified human cells that exceed 0.1 expected deaths per year would require specific international authorization to proceed. This standard would align the life sciences with quantitative safety expectations established across other major-hazard industries.

### SSL classifications for representative research scenarios

We applied the SSL framework using the tiered thresholds shown in Figure 2, where each SSL level spans an order of magnitude in expected harm. Research scenarios fall into two categories: agents that can be confidently classified using available transmission data, and agents requiring probabilistic assessment due to data limitations or uncertainty.

Using our transmission assessment (Methods), most BSL-4 agents are confidently classified as subcritical (Figure 4). Non-respiratory families like Filoviridae, Arenaviridae, and Nairoviridae default to subcritical. Ebola is SSL-3 based on observed IFR and historical outbreak sizes.

**Figure 4.**
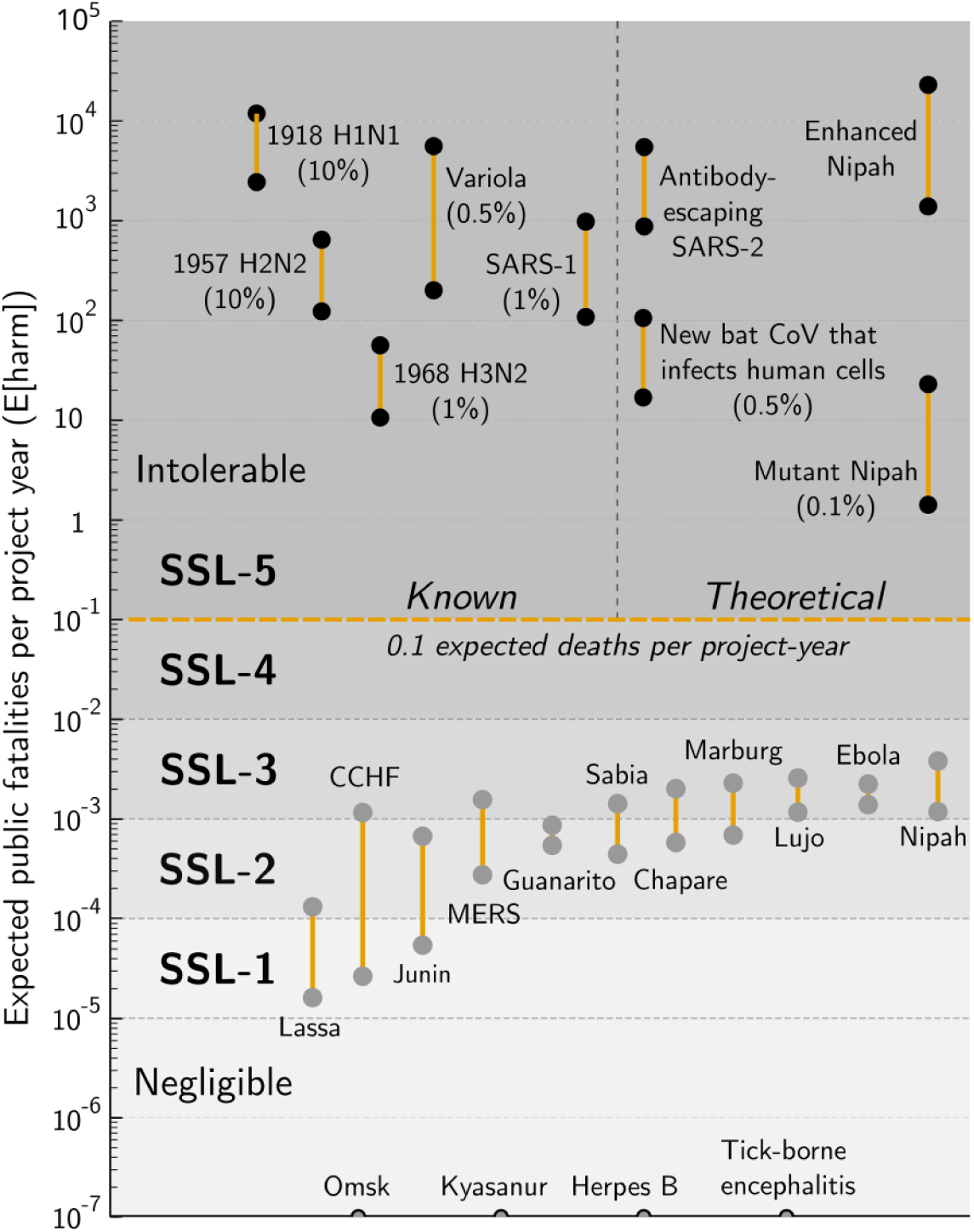
**| Societal Safety Levels for known and theoretical pathogens**. Expected harm from research involving potential pandemic pathogens and selected BSL-4 agents, assuming a lab-to-community release rate of 0.0025 per year (low-end Federal Select Agent Program assumed project-level baseline based on undetected and unreported exposures for registered entities; the NIH BSL3/4 baseline is up to 11-fold higher). Amplification probabilities are computed for one community seed using literature ranges for transmissibility (R) and overdispersion (k); total fatality C bands are anchored to historical estimates (Supplementary Information). Whiskers reflect conditional sensitivity from these R, k, and C ranges; scenario priors and the release rate are fixed per row. Percent values under each point denote *π* _super_, the fixed prior for the scenario: likelihood of population-immunity failure (influenza), export to LMIC settings before control (variola, SARS-1), or probability of pandemic capability (bat coronavirus computationally predicted to infect human cells; mutant Nipah). There is insufficient human evidence to classify Hendra virus (Methods). Exact parameter values are provided in the Supplementary Information.

Respiratory agents with extensive spillover evidence also qualify as subcritical, including Nipah virus (183+ documented spillovers without sustained transmission, SSL-3) and the BSL-3 MERS-CoV (2,500+ cases across multiple countries with limited human-to-human transmission, SSL-3). Effectively non-transmissible agents like tick-borne encephalitis and Kyasanur are SSL-1. Their low lethality relative to many BSL-3 agents raises separate questions about whether current BSL-4 classifications optimally reflect worker risk.

Not all agents can be confidently classified using available transmission data. Novel zoonotic viruses with limited spillover evidence, historical pandemic strains with uncertain population immunity, and products of pathogen enhancement research require regulatory decisions despite incomplete information. Explicit probabilistic assessment that acknowledges uncertainty is preferable to implicit judgment calls that obscure reasoning.

For these scenarios, we can establish the maximum tolerable probability that an agent exhibits sustained transmission (*π* _super,max_) by rearranging the expected harm equation:

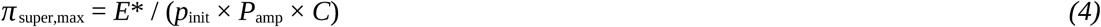

where *E\** = 0.1 deaths per year defines SSL-5. However, for policy clarity, we focus on the direct calculation: given conservative estimates of *π*_super_ based on available evidence, what is the expected harm? Table 1 shows these calculations for major research categories.

**Table 1:** Probabilistic assessment of known pathogens and theoretical experiments

| Experiment | $p_{\text{init}}$ | $\pi_{\text{super}}$ | $P_{\text{amp}}$ | $C$ | $E[\text{harm}]$ | SSL | Relevant facts |
| --- | --- | --- | --- | --- | --- | --- | --- |
| Hendra virus | 0.0025 | 0.005 | 0.07 | 10 million | 8.8 | 5 | Unknown $\pi_{\text{super}}$; 7 spillovers |
| Newly isolated bat coronavirus | 0.0025 | 0.005 | 0.22 | 27 million | 74.25 | 5 | 2 supercritical bat CoVs out of hundreds predicted to infect humans (~3,400 total) |
| 1918 H1N1 influenza | 0.0025 | 0.1 | 0.60 | 50 million | 7,500 | 5 | Population immunity, antibiotics = uncertainty<br>Antiviral prophylaxis may further reduce $p_{\text{init}}$ |
| Predicted vaccine-escaping SARS-CoV-2 | 0.0025 | 0.001 | 0.35 | 1 million | 0.875 | 5 | 1 in 1000 predictive success; would evade antibody but not T cell immunity |
| Successful enhanced transmission: H5N1 flu (GoF) | 0.0025 | 1 | 0.50 | 700,000 | 875 | 5 | Assumes lethality equivalent to 1977 H1N1: virulence- transmission tradeoff<br>Antiviral prophylaxis may further reduce $p_{\text{init}}$ |

### Agents with limited human transmission data

Hendra virus, the only BSL-4 agent from a high-risk viral family with fewer than 60 documented spillovers, lacks sufficient evidence to confidently rule out pandemic potential (Table 1).

Using conservative transmission parameters from its viral family (Paramyxoviridae), expected harm reaches 8.8 deaths per year – nearly 100-fold above the SSL-5 threshold. When assessing newly discovered zoonotic viruses from high-risk families, *π* _super_ depends on the strength of evidence for human transmissibility. Agents computationally predicted to infect human cells are lower-risk than agents isolated from infected humans, which are lower than agents with documented human-to-human transmission[42–45]. For example, newly isolated bat coronaviruses from the family that produced SARS-CoV-2 pose substantial risk. With at least two demonstrably pandemic-capable bat coronaviruses out of the hundreds predicted to infect human cells (from an estimated 3,400 total bat CoVs in nature[46]), even a conservative *π*_super_ estimate of 0.5% yields expected harm exceeding 70 deaths per year, over 700-fold above the threshold. To reach SSL-4, *π* _super_ would have to be less than one in ∼150,000 (0.0007%).

### Historical pandemic strains

For 1918 H1N1 influenza, *π*_super_ represents the probability that population immunity has waned enough to permit sustained transmission. While some cross-protection from descendant strains likely persists, and large antibiotic stockpiles (including veterinary) would reduce mortality from secondary bacterial infections, the historical consequences were so severe that even a 10% probability of renewed spread yields expected harm 75,000-fold above the SSL-5 threshold. In contrast, lab-adapted influenza strains such as H1N1 WSN33 and PR8 are deemed effectively subcritical due to attenuating mutations, as are strains isolated within the last 25 years due to population immunity.

### Enhancement experiments

Researchers creating mutations in endemic pathogens may inadvertently generate supercritical agents. Deep mutational scanning experiments creating single or double mutations pose minimal risk (*π* _super_ ≈ 0), as natural evolution samples such variants routinely. However, experiments testing combinatorial effects or especially AI-enabled redesigns of immune-targeted surface proteins in live-virus systems pose substantial risk. For SARS-CoV-2 spike variants designed to escape antibody immunity, even a 0.1% success rate yields expected harm approaching 1 death per year. The same experiment conducted using a pseudovirus system would pose negligible accidental societal risk because the system cannot transmit the target pathogen[47].

Experiments deliberately attempting to enhance the transmissibility of a subcritical pathogen[2,3] create the greatest risk. Because these studies aim to generate variants capable of widespread human transmission, successful experiments yield *π* _super_ = 1 by design. Enhanced H5N1 influenza, assuming substantially reduced lethality consistent with the 1977 H1N1 epidemic, would generate expected harm nearly 9,000-fold above the SSL-5 threshold. Any nontrivial probability of creating a transmissible variant (*π*_super_ × *P*_amp_ > 0.01) exceeds acceptable risk levels, effectively requiring international approval for work with replication-competent agents to proceed under the SSL framework.

### Molecular safeguards can reduce SSL ratings

The calculations above assume *p*_init_ = 0.0025 per year, reflecting undetected and unreported accidents that likely resulted in community exposure to live, replication-competent pathogens due to research performed by entities working with select agents[13]. Modern biotechnology provides multiple approaches that can substantially reduce this risk (Table 2).

**Table 2:**
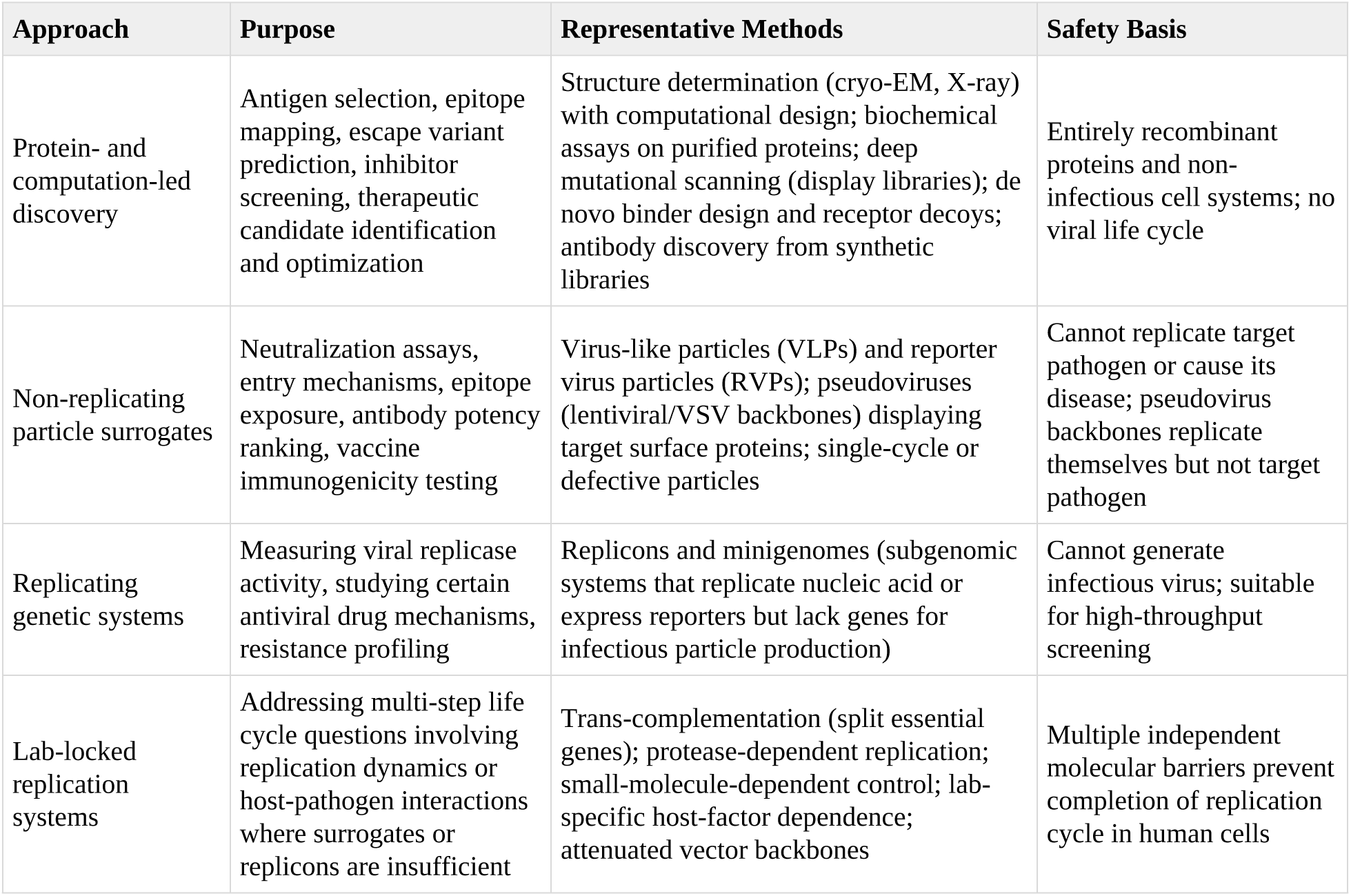
Methods of developing medical countermeasures without replication-competent virus

| Approach | Purpose | Representative Methods | Safety Basis |
| --- | --- | --- | --- |
| Protein- and computation-led discovery | Antigen selection, epitope mapping, escape variant prediction, inhibitor screening, therapeutic candidate identification and optimization | Structure determination (cryo-EM, X-ray) with computational design; biochemical assays on purified proteins; deep mutational scanning (display libraries); de novo binder design and receptor decoys; antibody discovery from synthetic libraries | Entirely recombinant proteins and non-infectious cell systems; no viral life cycle |
| Non-replicating particle surrogates | Neutralization assays, entry mechanisms, epitope exposure, antibody potency ranking, vaccine immunogenicity testing | Virus-like particles (VLPs) and reporter virus particles (RVPs); pseudoviruses (lentiviral/VSV backbones) displaying target surface proteins; single-cycle or defective particles | Cannot replicate target pathogen or cause its disease; pseudovirus backbones replicate themselves but not target pathogen |
| Replicating genetic systems | Measuring viral replicase activity, studying certain antiviral drug mechanisms, resistance profiling | Replicons and minigenomes (subgenomic systems that replicate nucleic acid or express reporters but lack genes for infectious particle production) | Cannot generate infectious virus; suitable for high-throughput screening |
| Lab-locked replication systems | Addressing multi-step life cycle questions involving replication dynamics or host-pathogen interactions where surrogates or replicons are insufficient | Trans-complementation (split essential genes); protease-dependent replication; small-molecule-dependent control; lab-specific host-factor dependence; attenuated vector backbones | Multiple independent molecular barriers prevent completion of replication cycle in human cells |

Computational models[48] and structural studies of pathogen proteins, including the prefusion-stabilized spike designs that enabled rapid COVID-19 vaccine development[49–52], pose zero transmission risk. Virus-like particles and pseudoviruses permit entry and neutralization studies but lack genes for replication[53,54]. Replicons replicate viral genetic material but cannot produce infectious particles[55]. Single-cycle packaging systems study the life cycle without sustained propagation[56]. At the highest-capability end, trans-complementation[57] (arguably including 6:2 reassortant influenza on an attenuated backbone[58]), small-molecule-dependent systems[59], unnatural amino acid requirements[60], and the insertion of human-specific miRNA targets[61] can recapitulate the sustained viral life cycle, but only in engineered laboratory environments featuring safe researcher-provided sequences or components absent from nature.

If two independent molecular barriers each provide 100-fold protection, expected harm could drop by four orders of magnitude, potentially shifting many SSL-5 experiments to SSL-2 or SSL-3. However, the magnitude of this risk reduction remains unknown: while the engineering principles are established and have been successfully applied to many viruses, barrier reliability under operational conditions has not yet been quantified. Until empirical studies demonstrate that specific molecular safeguards for viruses capable of sustained replication yield measured reductions in *p*_init_, regulators should assume baseline wild-type risk.

Quantitative validation of these methods could transform the regulatory landscape: the SSL framework would incentivize adoption of containment systems that achieve research objectives while maintaining public safety, while additionally providing an international approval pathway analogous to smallpox. This approach channels scientific effort toward methods that reduce societal risk rather than simply restricting access to dangerous agents, enabling high-consequence research to proceed with appropriate safeguards.

## Discussion

Current biosafety regulations have succeeded at protecting laboratory workers, with fewer than a hundred documented deaths from laboratory-acquired infections since biosafety began. Modern-day risks are comparable to or lower than most industrial occupations. Yet this success obscures ongoing neglect of societal risks by existing biosafety classifications.

The accumulated harm from all laboratory accidents represents a rounding error compared to a single pandemic. Current BSL ratings meticulously prevent harm to laboratory professionals without quantitatively evaluating risks to the billions of non-consenting members of the public. The Societal Safety Level framework addresses this gap by quantifying expected harm to society and establishing clear mitigation and approval thresholds. Our analysis reveals that studies of historical pandemic agents, transmission enhancement research on respiratory human pathogens, and laboratory characterization of certain live zoonotic viruses from high-risk families exceed the tolerable risk levels established across diverse industries.

### Policy implications

The SSL framework does not prohibit any research with potential pandemic pathogens, instead requiring specific approval for pathogens posing substantial public risks, in line with current restrictions on smallpox. Molecular containment systems can reduce public risk by orders of magnitude while preserving research capabilities (Figure 5). This creates appropriate incentives: researchers can pursue scientific questions using methods that achieve their objectives while maintaining public safety. The burden of proof shifts to those advocating wild-type replication-competent work. They must explain why safer alternatives are inadequate, and demonstrate through quantitative analysis that concrete expected benefits exceed quantified harms by a substantial margin. To date, no such analysis has been reported. Unlike medical advances with projected quantifiable benefits that exceed expected harms, abstract scientific knowledge and unspecified future benefits do not justify concrete risks the public would find intolerable in other contexts. SSL calculations could consequently provide the quantitative accidental-risk component of reviews required under the new U.S. policy; deliberate misuse and information hazards would require separate analysis[14].

**Figure 5.**
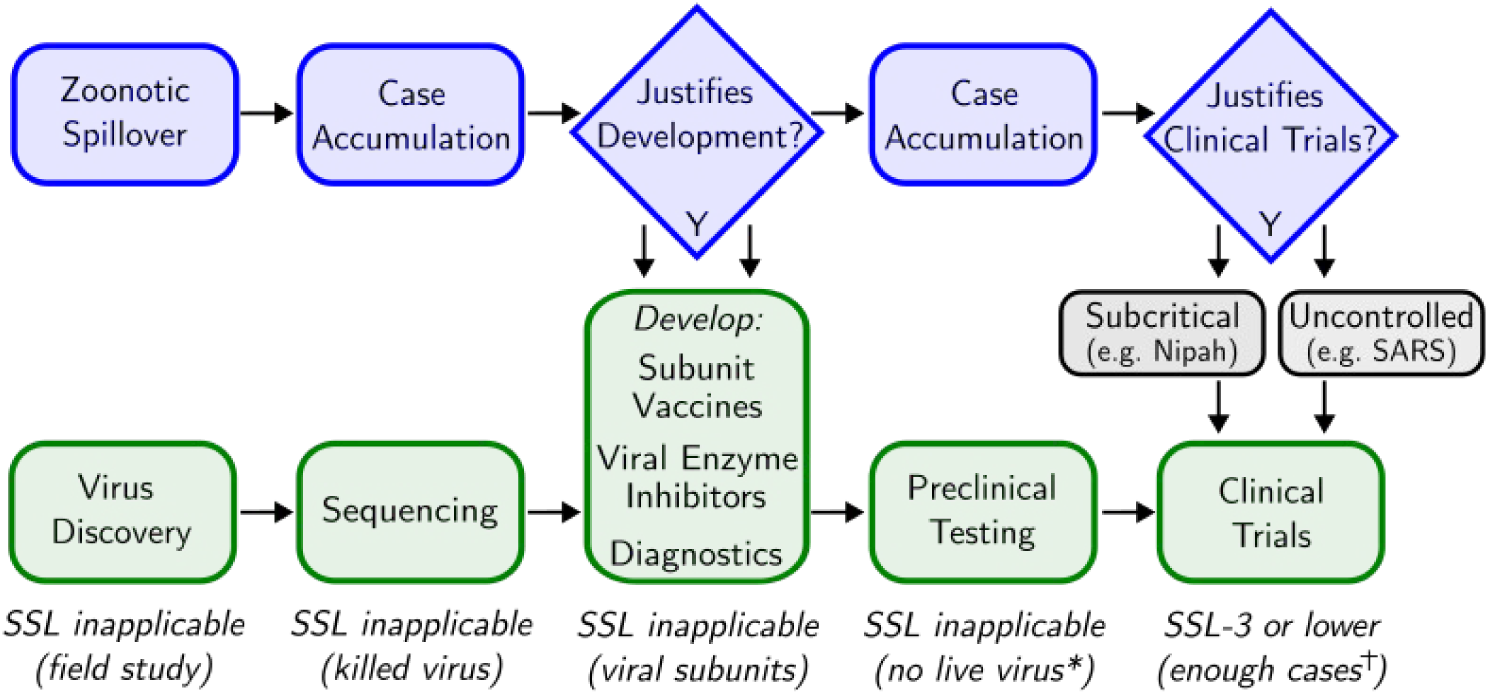
| Societal Safety Levels need not delay countermeasure development. Natural zoonotic spillover events (blue pathway) drive case accumulation that determines economic viability for countermeasure development and clinical trials. Research activities (green pathway) proceed independently using methods that do not require replication-competent pathogens. SSL classifications (bottom) show that restrictions apply only to high-risk laboratory work, not practical preparedness. *Preclinical testing uses pseudoviruses for inhibition studies and species-restricted chimeric backbones for animal challenge models, which do not pose transmission risks. †By the time sufficient cases accumulate to justify clinical trial investment, epidemiological evidence will have either demonstrated subcritical transmission (enabling SSL reclassification) or uncontrolled spread (making additional laboratory risk negligible). Mechanistic studies contributing to human knowledge can proceed on pathogens that do not create transmission risks for the general public.

For countermeasure development specifically, the safeguards detailed in Table 2 enable the full development pipeline from antigen discovery through preclinical validation without requiring wild-type replication-competent pathogens. Insofar as some regulators currently require wild-type virus for preclinical animal testing against seasonal influenza and other endemic human pathogens, there is no evident justification to apply similar requirements to potential pandemic pathogens, especially in light of expected-harm calculations.

The strongest potential case for wild-type experiments concerns historical pandemic strains extinct from human circulation: if these methods cannot validate countermeasures, and clinical trials are impossible due to eradication, then laboratory testing might appear necessary. The U.S. government previously determined that limited live-variola testing is necessary. However, most Table 2 techniques did not exist when those determinations were made. Even if the assessment still holds, variola is unique in lacking any closely related circulating disease for testing countermeasure efficacy (Supplementary Information, Box 1). Contemporary influenza strains provide adequate clinical validation pathways for historical influenza vaccines and therapeutics, eliminating any need for wild-type virus research.

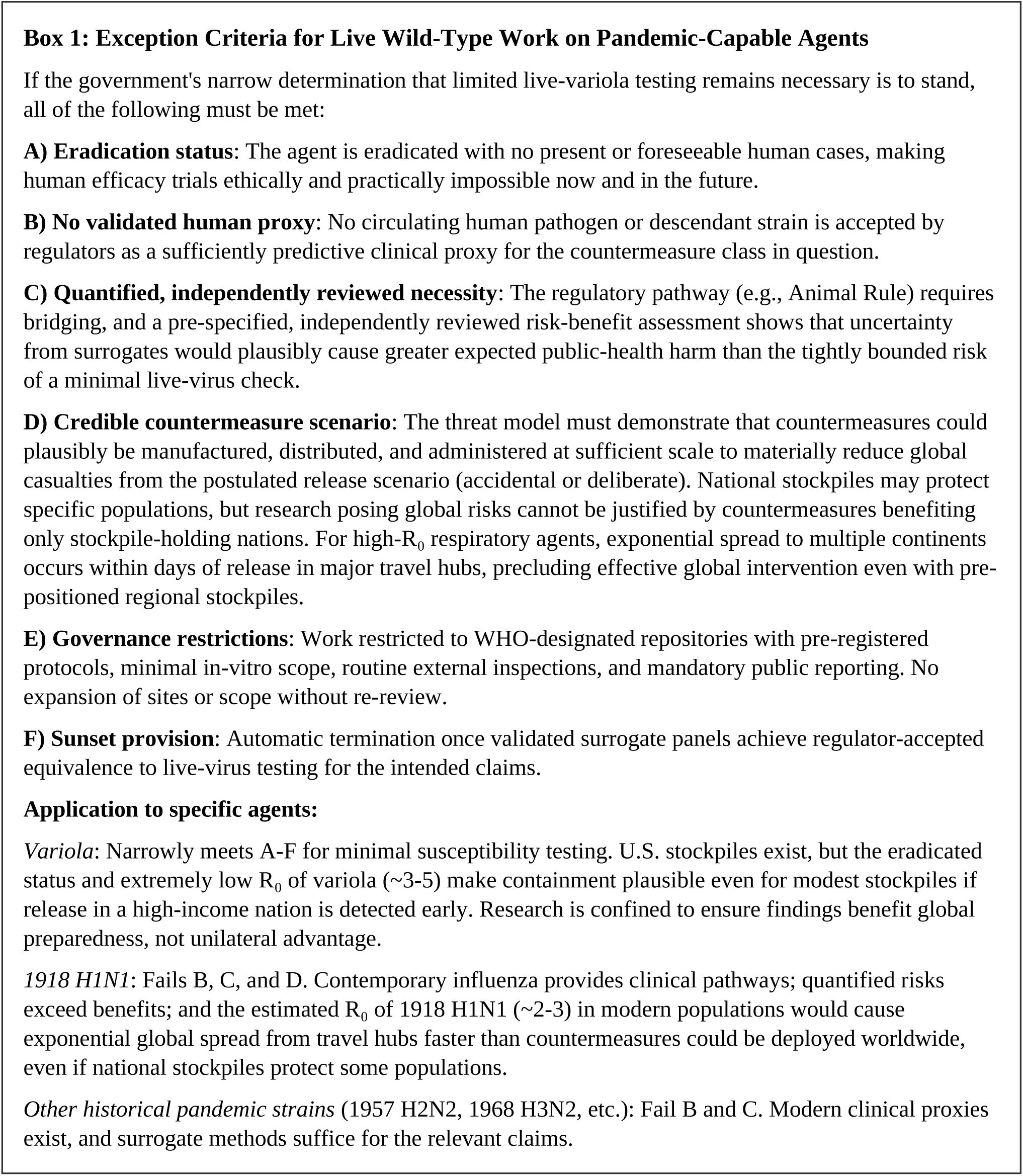

For naturally-occurring pathogens still causing human infections, laboratory experiments with wild-type replication-competent agents face a high burden of justification (Fig. 5). Initial spillovers with few cases do not warrant countermeasure investment. Once sufficient infections accumulate to justify development, the techniques in Table 2 enable preclinical work. By the time cases accumulate to justify validation costs, epidemiological evidence typically reveals one of two outcomes: either repeated spillovers without sustained transmission demonstrate subcriticality (permitting SSL-1/2/3 work, as with Nipah), or pandemic spread enables field validation (as with SARS-CoV-2).

The available evidence supports this framework. Discovering and characterizing viruses in animals before outbreaks has not historically accelerated countermeasure development; most priority pathogens were identified during human infections. Quantitative modeling suggests community-based monitoring at the animal-human interface may offer greater benefits than individual virus characterization, though additional research could refine these estimates.

Importantly, the quantitative nature of the SSL framework is bidirectional: it demonstrates that many agents currently require higher levels of containment than is warranted. By permitting BSLs to be assigned independent of transmissibility, the adoption of SSLs should trigger the reevaluation and potentially reclassification of all non-transmissible BSL-4 agents as well as certain subcritical agents. For example, there is little justification for Omsk hemorrhagic fever, which is non-transmissible (SSL-1) and seldom lethal, to require BSL-4 when the far more lethal and subcritically transmissible (SSL-3) MERS-CoV requires BSL-3. The biosafety ratchet, whereby requirements accumulate but rarely decrease, misaligns resources and scientific incentives. Explicit de-escalation criteria are a feature of the quantitative framework, not a limitation.

### Implementation

Adopting the SSL framework requires three steps. First, researchers calculate expected harm for their proposed work using observed accident rates, transmission parameters from the literature, and consequence estimates. Second, institutional biosafety committees and federal oversight bodies review these calculations and adjust for locally relevant factors. Third, projects exceeding SSL thresholds must either adopt safer methods (molecular containment systems, non-replicating surrogates), demonstrate the agent is actually subcritical through epidemiological evidence (Methods), or obtain explicit permission from an authorized international body after public debate. Novel agents with limited human case data are treated as high-risk until evidence demonstrates otherwise. For projects reviewed as potential dangerous gain-of-function research under the 2026 policy, the SSL estimate could supply the quantitative accidental-harm component of the required risk-benefit assessment[14].

The framework incentivizes continuous safety improvement, but realizing this potential requires robust incident data and adequate resources. Current reporting systems may create disincentives for disclosure; the aviation industry addressed a similar challenge through the Aviation Safety Reporting System, which provides immunity from punitive action for self-reported incidents and has dramatically improved safety by enabling systematic learning from near-misses. An analogous non-punitive framework for laboratory incidents, ideally international in scope, would enable facilities to benchmark performance, identify best practices, and surface leading indicators such as training gaps and procedural failures. Quantitative thresholds also clarify resource needs: many institutions struggle with aging facilities and understaffed biosafety programs, and SSL classifications would help justify funding commensurate with actual risk.

Institutional biosafety committees reviewing SSL-3 and above should document explicit risk-benefit reasoning, including quantified expected harms and concrete anticipated benefits. For SSL-4 and SSL-5 work, this documentation would inform review by national oversight bodies and, where required, international authorization processes. Where committees lack expertise in ethical risk assessment, additional resources should be provided for external consultation. These expectations parallel existing requirements for human subjects research and would formalize deliberation that currently occurs informally or not at all.

Notably, SSLs would impose few if any additional requirements on work already conducted at BSL-4, much of which might reasonably be reduced to BSL-3. Virtually all of the impact would apply to a narrow category of research involving pandemic-capable agents at lower containment levels, chiefly novel bat coronaviruses, historical pandemic influenza strains, enhanced-transmission experiments, and to a lesser extent, MERS-CoV. As containment methods improve and incident rates decline, expected harm calculations automatically adjust, potentially permitting work that previously exceeded thresholds. This creates the right incentive structure: researchers gain access to high-consequence agents by demonstrating superior safety measures.

The public tolerates chemical plants and nuclear facilities near their communities not because they trust these industries, but because strict quantitative limits constrain the risks these operations may impose. The life sciences have no equivalent framework for pandemic-capable pathogens. Adopting SSLs would establish the same enforceable risk thresholds, bringing high-consequence biological research into alignment with the quantitative safety standards that allow other hazardous but beneficial activities to operate with public acceptance.

## Methods

### Framework overview

We quantify societal risk as expected deaths per year, including laboratory workers. Our analysis considers three pathogen classes: subcritical agents (*R*_eff_ < 1, cannot sustain transmission or are already present), supercritical agents (*R*_eff_ > 1, can cause pandemics), and agents with uncertain transmission potential. Expected harm incorporates: (i) probability of laboratory release reaching the community (*p*_init_), (ii) probability a release amplifies into sustained spread (*P*_amp_), and (iii) deaths if amplification occurs (C).

### Parameter estimation and sources

Transmission parameters (R, k) were drawn from published estimates for well-characterized agents; for novel agents lacking direct estimates, we assigned conservative values from the most transmissible characterized member of the same viral family. Infection fatality rates derive from outbreak investigations and seroprevalence studies. Consequence estimates (C) are anchored to historical pandemic death tolls, scaled for modern population size, healthcare capacity, and residual immunity where applicable. Supplementary Tables S1-S7 provide complete outbreak totals, parameter values, ranges, and primary sources for all agents and scenarios analyzed. Where multiple credible estimates exist, we report ranges; where estimates are contested, we use values that yield conservative (lower) expected harm calculations.

### Laboratory-acquired infection rates

We estimated modern BSL-4 worker fatality risk using the single confirmed death (Ebola, VECTOR Institute, 2004) and global workforce estimates. With ∼50 operating BSL-4 facilities employing 3,000-10,000 authorized personnel, the fatal infection rate is 0.3-6 per 100,000 worker-years—comparable to most industrial occupations. Historical incidents (Marburg 1967, Birmingham smallpox 1978, Yale Lassa 1969) and Herpes B occupational deaths from animal facilities are tracked separately.

### Release probability (*p*_init_)

We estimated community release rates from U.S. laboratory incident data. The Federal Select Agent Program (2003-2017) recorded 10 community-exposed infections across 4,067 entity-years, yielding *p*_init_ = 0.0025 per entity year. Entities may have multiple projects with a single agent, actively study multiple agents, or work with registered agent for less than a full year; in the absence of more reliable data, we assume entity-years are equivalent to project-years with a relevant detectable agent. NIH oversight records (2004-2017) suggest higher rates of 0.016-0.028 per year. We use 0.0025 as baseline for conservative risk assessment; programs with N simultaneous projects over Y years have cumulative release probability 1-(1-*p*_init_)^NY. Containment improvements reduce *p*_init_ proportionally.

### Classifying transmission potential

Most agents can be classified using available evidence without requiring extensive human transmission studies:

*Agents from low-risk viral families* (Filoviridae, Arenaviridae, Nairoviridae, mosquito or tick-borne Flaviviridae): These families exhibit limited respiratory human-to-human transmission based on historical outbreak patterns and transmission routes. Default to subcritical unless evidence shows efficient respiratory transmission.
*Agents from high-risk families* (Orthomyxoviridae, Coronaviridae, Paramyxoviridae, Orthopoxviridae): Require spillover evidence to demonstrate subcriticality. We apply rule-of-3 statistics: if M independent spillovers with community exposure failed to cause sustained transmission, pandemic probability per episode ≤ 3/M (95% confidence)[62]. For M ≥ 60, we presume subcriticality. For M < 60, insufficient evidence—agent remains high-risk pending additional data.
*Example applications:* Nipah virus (Paramyxoviridae) has 183+ documented spillovers without sustained transmission, qualifying as subcritical despite high-risk family membership. Ebola and Lassa default to subcritical based on family and transmission route. Hendra (Paramyxoviridae) has only 7 human cases— insufficient evidence for confident classification. Variola is supercritical based on historical sustained transmission.

### Amplification probability (P_amp_)

For supercritical agents, we model early community spread as a branching process with negative binomial offspring distribution (mean *R*, dispersion *k*). The extinction probability q solves q = [p/(1-(1-p)q)]^*k* where p = *k*/(*k*+*R*), computed iteratively to 10^-8 tolerance. Amplification probability is *P*_amp_ = 1-q.

We assigned family-specific parameters from literature: influenza *R* = 1.4-2.4, *k* = 1-2 (minimal overdispersion); SARS-class coronaviruses *R* = 2-3, k = 0.1-0.3 (strong overdispersion); SARS-CoV-2 omicron variants *R* = 6-8, k = 0.1-0.2; variola *R* = 3-6, k = 0.5-1.0. Higher *R* and *k* both increase *P*_amp_. For *R* = 2.5 with *k* = 0.2, *P*_amp_ ≈ 0.244; with *k* = 2, *P*_amp_ ≈ 0.725. Our behavioral response model assumes rapid recognition of novel pathogens. For agents with mild or influenza-like presentation, recognition may be delayed by multiple transmission generations, during which behavioral modification is minimal. Our estimates may therefore be conservative for the pathogen category of greatest concern: transmissible agents with nonspecific symptoms.

### Consequence estimation (*C*)

For novel pandemics, we anchored *C* to historical precedents: 1918-class influenza 20-80 million deaths (adjusted for modern antibiotics and healthcare); 1957/1968 pandemics ∼2 million; SARS-CoV-2 ∼27 million. When a closely related agent is already spreading but has not yet infected most people, we calculate incremental deaths – the additional burden from the supercritical laboratory strain racing ahead of wild-type spread. For highly lethal agents (variola, enhanced Nipah), we used ranges reflecting severity-dependent behavioral response and countermeasure availability.

### Subcritical agents

When *R*_eff_ < 1, releases cause limited chains that self-extinguish. Expected deaths are *p*_init_ × IFR/(1-*R*_eff_), where IFR is infection fatality rate. We modeled *R*_eff_ = *R* × [c + (1-c)(1-b)] where c is the fraction of covert transmission (pre-symptomatic/asymptomatic), and b is behavioral contact reduction triggered by symptom severity; parameters were estimated from literature on transmission timing and hospitalization or fatality rates. Highly lethal agents with distinctive symptoms (Ebola, SARS-CoV-1) have low c and high b, yielding *R*_eff_ well below intrinsic R. Mild or influenza-like illnesses have higher c, limiting behavioral impact.

### Agents with uncertain transmission potential

For novel zoonotic agents, historical pandemics with waning immunity, and enhancement experiments, we use π_super_ (probability of supercritical behavior) to handle uncertainty:

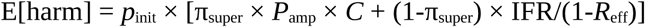

When supercritical consequences dominate, this simplifies to *p*_init_ × π_super_ × *P*_amp_ × *C*.

For novel zoonoses, we bounded π_super_ using spillover evidence and prediction reliability. If M similar viruses spilled over without pandemic, π_super_ ≤ 3/M. For computationally predicted viruses, we adjusted for low base rates and predictor specificity—even high-accuracy classifiers produce many false positives when true prevalence is low. For viruses isolated from infected humans with K failed transmission chains, π_super_ ≤ 3/K when amplification probability given supercriticality exceeds the observed chain-extinction rate.

For historical pandemic strains, π_super_ represents probability that population immunity has waned sufficiently for *R*_eff_ > 1. For 1918 H1N1, we estimated π_super_ = 0.1 based on genetic distance from circulating strains and cohort immunity data. Laboratory-adapted strains (WSN33, PR8/34) were assigned π_super_ ≈ 0 due to attenuating mutations.

For enhancement experiments, π_super_ reflects success probability. Deep mutational scanning (single/double mutants) has π_super_ ≈ 0 because natural evolution samples such variants. Combinatorial enhancement or AI-enabled immune escape design targeting surfaces exceeding natural variation has substantial π_super_. Deliberate transmission-enhancement experiments have π_super_ = 1 by design if successful.

### SSL threshold determination

Multiple industries converge on ∼0.1 expected deaths per facility-year: Australian dam safety (10^-6 individual risk × 100,000 downstream), UK chemical facilities (10^-6 × ∼100,000 within impact radius), NRC nuclear safety goals (10^-6 × 10,000-100,000), and regional airport safety (observed ∼5×10^-6 per flight × 20,000 annual flights). We adopted E* = 0.1 deaths per project- year as the SSL-5 threshold. A more comprehensive compilation of industry sources is available in the Supplementary Information.

This threshold enables direct calculation of maximum tolerable values:

● Maximum π_super_ = 0.1/(*p*_init_ × *P*_amp_ × *C*)
● Maximum *p*_init_ = 0.1/(π_super_ × *P*_amp_ × *C*)
● Maximum *P*_amp_ = 0.1/(*p*_init_ × π_super_ × *C*)

These expressions identify whether compliance requires improved containment (reduce *p*_init_), transmission blocking (reduce *P*_amp_ via countermeasures), or consequence mitigation (reduce *C* via vaccines/therapeutics).

### Uncertainty and presentation

We computed expected harm ranges by varying *R*, *k*, and *C* within literature-supported bands. All calculations assume single-seed releases; we do not model simultaneous multi-laboratory incidents. To avoid false precision and limit misuse, we display *P*_amp_ and *E*[harm] to two significant figures, with π_super_ × *P*_amp_ capped at 10% in public figures (except variola). All risk estimates involve uncertainty. The alternative to imprecise quantitative assessment is not more precise assessment, but no quantitative assessment at all.

## Author Contributions

K.M.E. conceived the study, developed the framework, performed the analyses, and wrote the initial draft. L.J. contributed to figure development and manuscript editing. D.G. contributed to manuscript editing. All authors reviewed and approved the final manuscript.

## Acknowledgements

We are grateful to numerous virologists and biosafety analysts for helpful feedback. This study was supported by a gift from Coefficient Giving (to K.M.E.) and a Draper Scholarship (to L.J.). The funders played no role in the conception, development, or decision to publish.

---

**Supplementary Figure S1.**
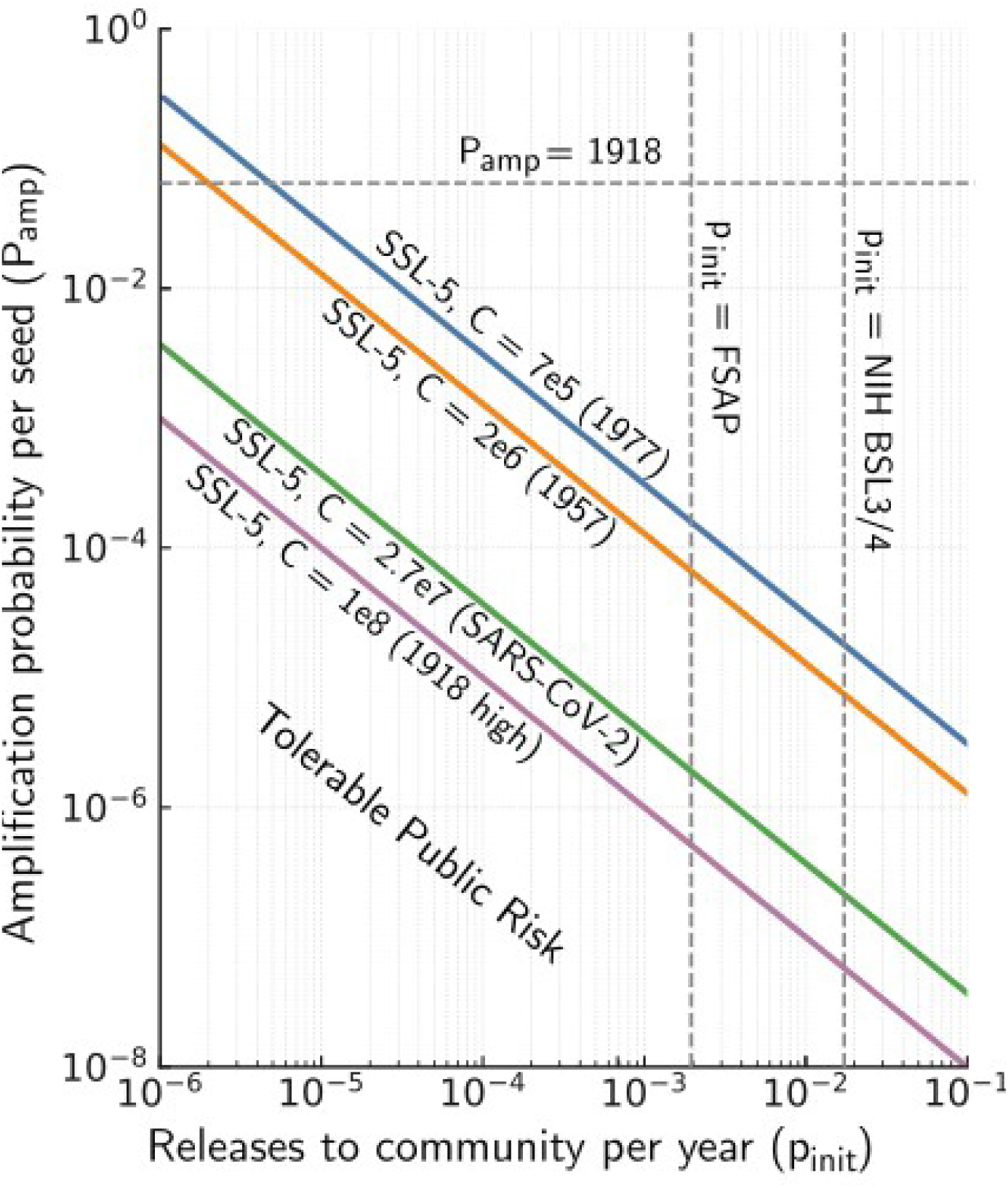
| SSL-5 boundaries depend on pandemic consequences. The SSL-5 threshold (0.1 expected deaths per year) creates hyperbolic boundaries in pinit-Pamp space for different consequence scales. Historical pandemic scenarios span five orders of magnitude in C, from 1977 H1N1 (C ≈ 700,000 estimated deaths) to worst-case 1918 H1N1 (C ≈ 100 million deaths). Vertical dashed lines show empirical accident rates from U.S. laboratories; horizontal lines show estimated amplification probabilities for SARS-CoV-2 and 1918 H1N1. Most historical pandemic agents fall above their respective SSL-5 boundaries at current containment performance, indicating that marked safety improvements or alternative methods (Table 2) are required.

**Supplementary Table S1.**
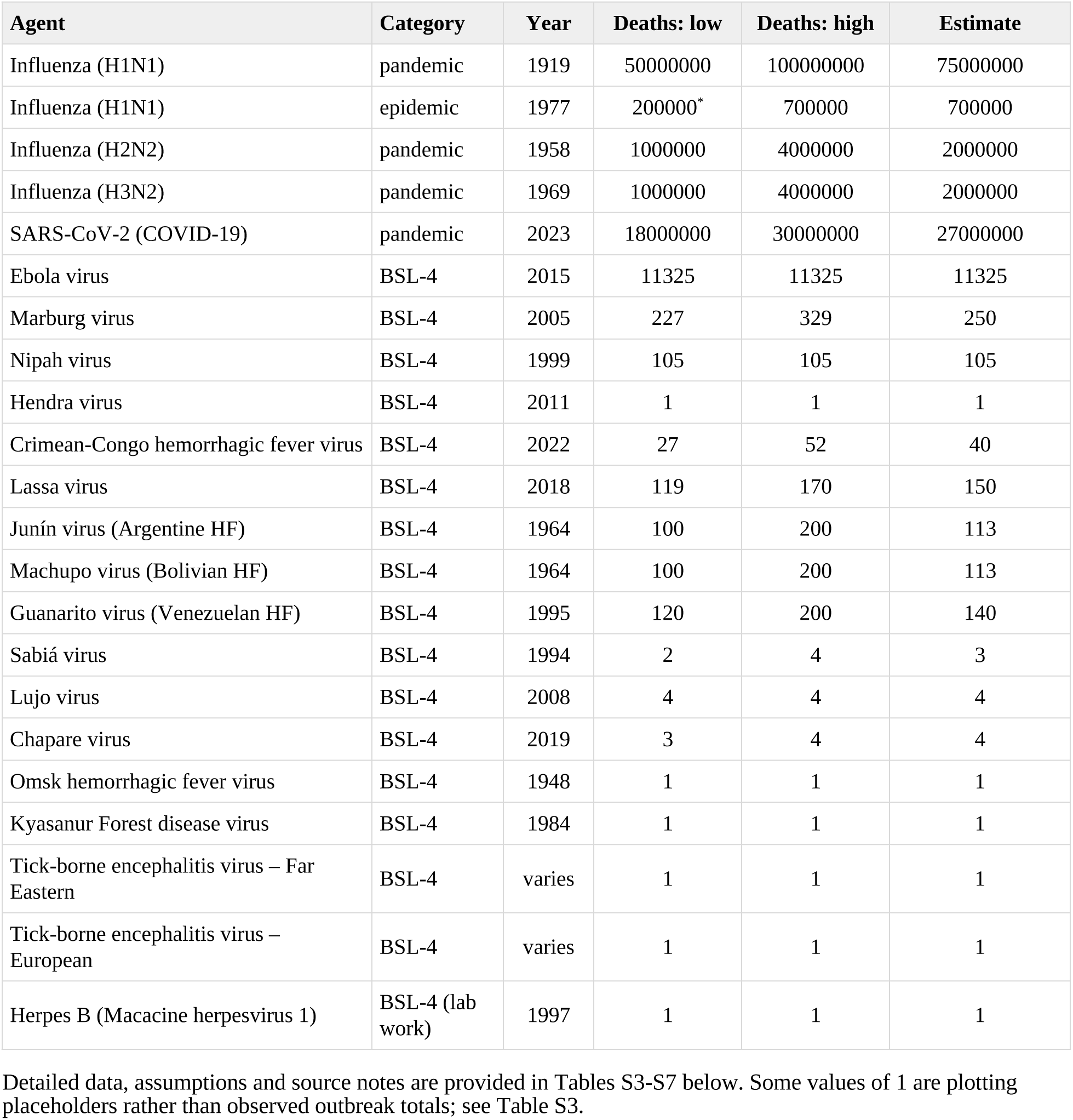
| Largest single natural outbreaks of all pathogens and of BSL-4 agents. Data compiled from referenced sources. *Assumes total influenza mortality was partially constrained by cross-reactive immunity or behavioral responses, such that the 1977 H1N1 emergence displaced deaths that would otherwise have occurred from seasonal H3N2.

**Supplementary Table S2.**
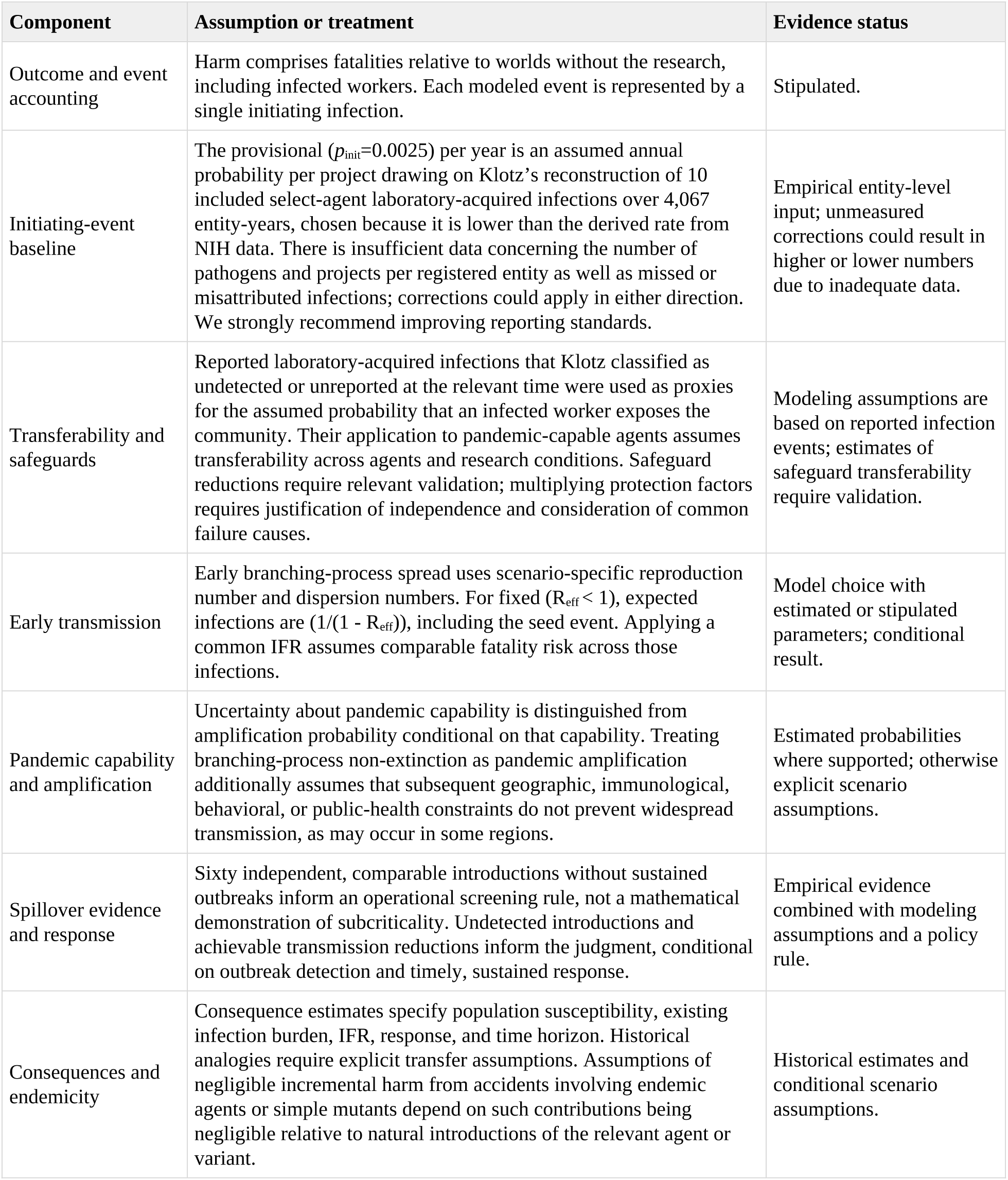
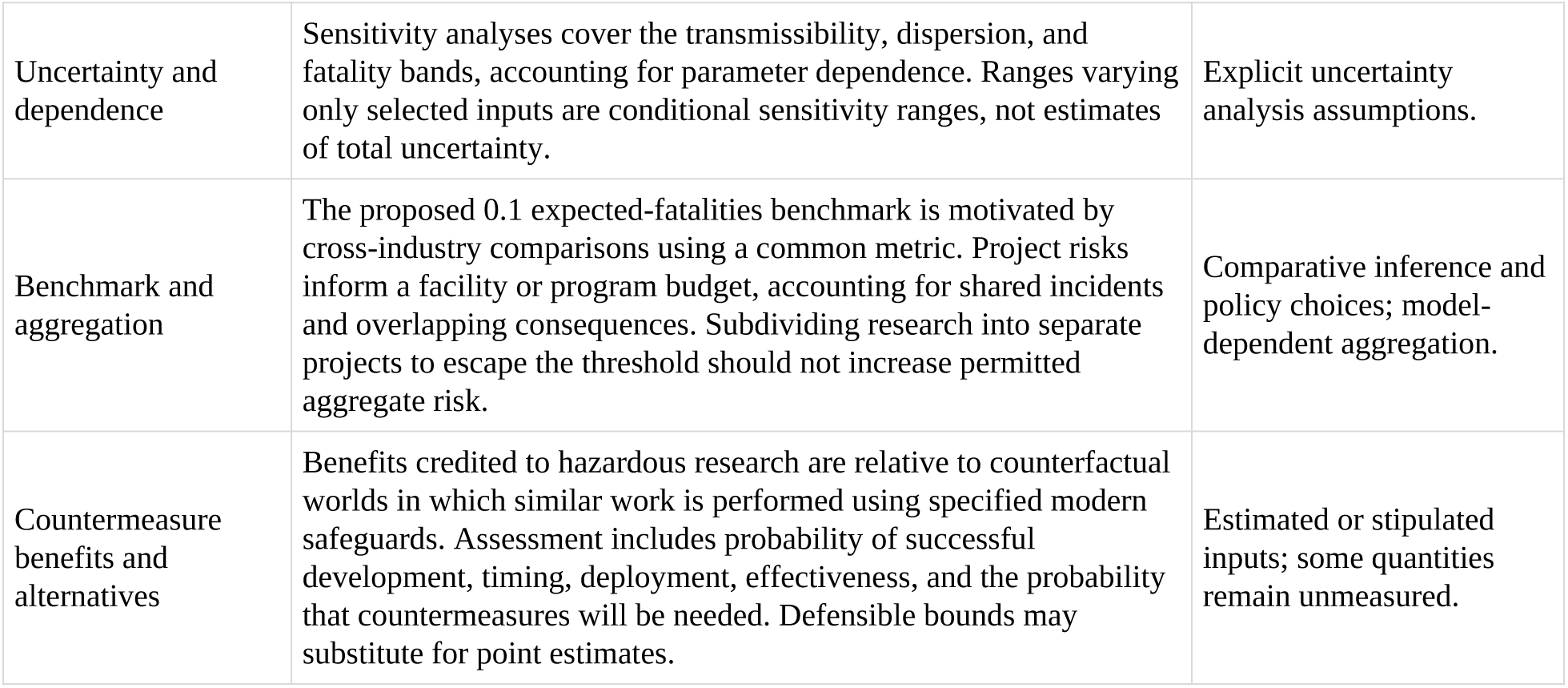
| Limitations and caveats. Parameter values and scenario ranges are based on available data and extrapolations thereof. To convey uncertainty, we stipulate explicit assumptions, sensitivity analyses, and evidence status. Improved incident reporting standards and quantifying escape rates and independence of laboratory safeguards can substantially reduce uncertainties regarding assessed risks and the associated SSL status of project proposals.

