## Supplementary Tables and Figures for "Complementing Biosafety Levels With Societal Safety Levels"

**Supplementary Table S1 | Largest single natural outbreaks of all pathogens and of BSL-4 agents.** Data compiled from referenced sources. *Assumes total influenza mortality was partially constrained by cross-reactive immunity or behavioral responses, such that the 1977 H1N1 emergence displaced deaths that would otherwise have occurred from seasonal H3N2.

| Agent | Category | Year | Deaths: low | Deaths: high | Estimate |
|---|---|---|---|---|---|
| Influenza (H1N1) | pandemic | 1919 | 50000000 | 100000000 | 75000000 |
| Influenza (H1N1) | epidemic | 1977 | 200000* | 700000 | 700000 |
| Influenza (H2N2) | pandemic | 1958 | 1000000 | 4000000 | 2000000 |
| Influenza (H3N2) | pandemic | 1969 | 1000000 | 4000000 | 2000000 |
| SARS-CoV-2 (COVID-19) | pandemic | 2023 | 18000000 | 30000000 | 27000000 |
| Ebola virus | BSL-4 | 2015 | 11325 | 11325 | 11325 |
| Marburg virus | BSL-4 | 2005 | 227 | 329 | 250 |
| Nipah virus | BSL-4 | 1999 | 105 | 105 | 105 |
| Hendra virus | BSL-4 | 2011 | 1 | 1 | 1 |
| Crimean-Congo hemorrhagic fever virus | BSL-4 | 2022 | 27 | 52 | 40 |
| Lassa virus | BSL-4 | 2018 | 119 | 170 | 150 |
| Junín virus (Argentine HF) | BSL-4 | 1964 | 100 | 200 | 113 |
| Machupo virus (Bolivian HF) | BSL-4 | 1964 | 100 | 200 | 113 |
| Guanarito virus (Venezuelan HF) | BSL-4 | 1995 | 120 | 200 | 140 |
| Sabiá virus | BSL-4 | 1994 | 2 | 4 | 3 |
| Lujo virus | BSL-4 | 2008 | 4 | 4 | 4 |
| Chapare virus | BSL-4 | 2019 | 3 | 4 | 4 |
| Omsk hemorrhagic fever virus | BSL-4 | 1948 | 1 | 1 | 1 |
| Kyasanur Forest disease virus | BSL-4 | 1984 | 1 | 1 | 1 |
| Tick-borne encephalitis virus – Far Eastern | BSL-4 | varies | 1 | 1 | 1 |
| Tick-borne encephalitis virus – European | BSL-4 | varies | 1 | 1 | 1 |
| Herpes B (Macacine herpesvirus 1) | BSL-4 (lab work) | 1997 | 1 | 1 | 1 |

Detailed data, assumptions and source notes are provided in Tables S3-S7 below. Some values of 1 are plotting placeholders rather than observed outbreak totals; see Table S3.

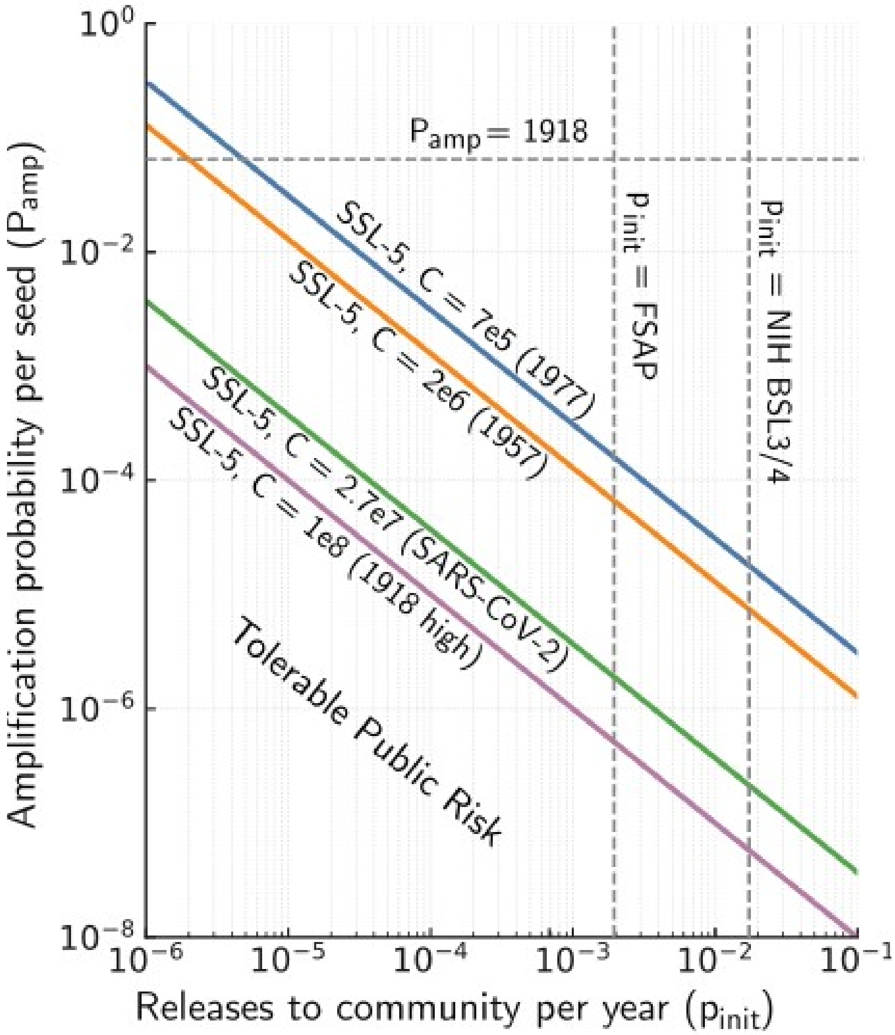


**Supplementary Figure S1 | SSL-5 boundaries depend on pandemic consequences.** The SSL-5 threshold (0.1 expected deaths per year) creates hyperbolic boundaries in pinit-Pamp space for different consequence scales. Historical pandemic scenarios span five orders of magnitude in C, from 1977 H1N1 (C ≈ 700,000 estimated deaths) to worst-case 1918 H1N1 (C ≈ 100 million deaths). Vertical dashed lines show empirical accident rates from U.S. laboratories; horizontal lines show estimated amplification probabilities for SARS-CoV-2 and 1918 H1N1. Most historical pandemic agents fall above their respective SSL-5 boundaries at current containment performance, indicating that marked safety improvements or alternative methods (Table 2) are required.

**Box 1: Exception Criteria for Live Wild-Type Work on Pandemic-Capable Agents**

If the government's narrow determination that limited live-variola testing remains necessary is to stand, all of the following must be met:

**A) Eradication status**: The agent is eradicated with no present or foreseeable human cases, making human efficacy trials ethically and practically impossible now and in the future.

**B) No validated human proxy**: No circulating human pathogen or descendant strain is accepted by regulators as a sufficiently predictive clinical proxy for the countermeasure class in question.

**C) Quantified, independently reviewed necessity**: The regulatory pathway (e.g., Animal Rule) requires bridging, and a pre-specified, independently reviewed risk-benefit assessment shows that uncertainty from surrogates would plausibly cause greater expected public-health harm than the tightly bounded risk of a minimal live-virus check.

**D) Credible countermeasure scenario**: The threat model must demonstrate that countermeasures could plausibly be manufactured, distributed, and administered at sufficient scale to materially reduce global casualties from the postulated release scenario (accidental or deliberate). National stockpiles may protect specific populations, but research posing global risks cannot be justified by countermeasures benefiting only stockpile-holding nations. For high-$R_0$ respiratory agents, exponential spread to multiple continents occurs within days of release in major travel hubs, precluding effective global intervention even with pre-positioned regional stockpiles.

**E) Governance restrictions**: Work restricted to WHO-designated repositories with pre-registered protocols, minimal in-vitro scope, routine external inspections, and mandatory public reporting. No expansion of sites or scope without re-review.

**F) Sunset provision**: Automatic termination once validated surrogate panels achieve regulator-accepted equivalence to live-virus testing for the intended claims.

**Application to specific agents:**

*Variola*: Narrowly meets A-F for minimal susceptibility testing. U.S. stockpiles exist, but the eradicated status and extremely low $R_0$ of variola (~3-5) make containment plausible even for modest stockpiles if release in a high-income nation is detected early. Research is confined to ensure findings benefit global preparedness, not unilateral advantage.

*1918 H1N1*: Fails B, C, and D. Contemporary influenza provides clinical pathways; quantified risks exceed benefits; and the estimated $R_0$ of 1918 H1N1 (~2-3) in modern populations would cause exponential global spread from travel hubs faster than countermeasures could be deployed worldwide, even if national stockpiles protect some populations.

*Other historical pandemic strains* (1957 H2N2, 1968 H3N2, etc.): Fail B and C. Modern clinical proxies exist, and surrogate methods suffice for the relevant claims.

**Supplementary Table S2 | Limitations and caveats.** Parameter values and scenario ranges are based on available data and extrapolations thereof. To convey uncertainty, we stipulate explicit assumptions, sensitivity analyses, and evidence status. Improved incident reporting standards and quantifying escape rates and independence of laboratory safeguards can substantially reduce uncertainties regarding assessed risks and the associated SSL status of project proposals.

| Component | Assumption or treatment | Evidence status |
|---|---|---|
| Outcome and event accounting | Harm comprises fatalities relative to worlds without the research, including infected workers. Each modeled event is represented by a single initiating infection. | Stipulated. |
| Initiating-event baseline | The provisional ($p_{init}$=0.0025) per year is an assumed annual probability per project drawing on Klotz's reconstruction of 10 included select-agent laboratory-acquired infections over 4,067 entity-years, chosen because it is lower than the derived rate from NIH data. There is insufficient data concerning the number of pathogens and projects per registered entity as well as missed or misattributed infections; corrections could apply in either direction. We strongly recommend improving reporting standards. | Empirical entity-level input; unmeasured corrections could result in higher or lower numbers due to inadequate data. |
| Transferability and safeguards | Reported laboratory-acquired infections that Klotz classified as undetected or unreported at the relevant time were used as proxies for the assumed probability that an infected worker exposes the community. Their application to pandemic-capable agents assumes transferability across agents and research conditions. Safeguard reductions require relevant validation; multiplying protection factors requires justification of independence and consideration of common failure causes. | Modeling assumptions are based on reported infection events; estimates of safeguard transferability require validation. |
| Early transmission | Early branching-process spread uses scenario-specific reproduction number and dispersion numbers. For fixed ($R_{eff} < 1$), expected infections are ($1/(1 - R_{eff})$), including the seed event. Applying a common IFR assumes comparable fatality risk across those infections. | Model choice with estimated or stipulated parameters; conditional result. |
| Pandemic capability and amplification | Uncertainty about pandemic capability is distinguished from amplification probability conditional on that capability. Treating branching-process non-extinction as pandemic amplification additionally assumes that subsequent geographic, immunological, behavioral, or public-health constraints do not prevent widespread transmission, as may occur in some regions. | Estimated probabilities where supported; otherwise explicit scenario assumptions. |
| Spillover evidence and response | Sixty independent, comparable introductions without sustained outbreaks inform an operational screening rule, not a mathematical demonstration of subcriticality. Undetected introductions and achievable transmission reductions inform the judgment, conditional on outbreak detection and timely, sustained response. | Empirical evidence combined with modeling assumptions and a policy rule. |
| Consequences and endemicity | Consequence estimates specify population susceptibility, existing infection burden, IFR, response, and time horizon. Historical analogies require explicit transfer assumptions. Assumptions of negligible incremental harm from accidents involving endemic agents or simple mutants depend on such contributions being negligible relative to natural introductions of the relevant agent or variant. | Historical estimates and conditional scenario assumptions. |

| Component | Assumption or treatment | Evidence status |
|---|---|---|
| Uncertainty and dependence | Sensitivity analyses cover the transmissibility, dispersion, and fatality bands, accounting for parameter dependence. Ranges varying only selected inputs are conditional sensitivity ranges, not estimates of total uncertainty. | Explicit uncertainty analysis assumptions. |
| Benchmark and aggregation | The proposed 0.1 expected-fatalities benchmark is motivated by cross-industry comparisons using a common metric. Project risks inform a facility or program budget, accounting for shared incidents and overlapping consequences. Subdividing research into separate projects to escape the threshold should not increase permitted aggregate risk. | Comparative inference and policy choices; model-dependent aggregation. |
| Countermeasure benefits and alternatives | Benefits credited to hazardous research are relative to counterfactual worlds in which similar work is performed using specified modern safeguards. Assessment includes probability of successful development, timing, deployment, effectiveness, and the probability that countermeasures will be needed. Defensible bounds may substitute for point estimates. | Estimated or stipulated inputs; some quantities remain unmeasured. |

Tables S3-S7 and their source references follow. The calculation workbook is linked in the accompanying calculation notes.

#### Supplementary Table S3. Outbreak benchmarks and source notes

Detailed source data corresponding to Table S1. Best estimates and low-high values are reproduced from the workbook. They include outbreak totals, cumulative estimates and explicitly identified plotting placeholders; these are distinguished in the notes.

| Agent | Category | Year | Best estimate | Low to high | Interpretation and sources |
|---|---|---|---|---|---|
| Influenza (H1N1) | pandemic | 1919 | 75,000,000 | 50,000,000 to 100,000,000 | Global estimate range [1] |
| Influenza (H1N1) | epidemic | 1977 | 700,000 | 200,000 to 700,000 | Lower estimate requires the unsupported assumption that seasonal H3N2 or swine H1N1 would have counterfactually caused as many deaths [2] |
| Influenza (H2N2) | pandemic | 1958 | 2,000,000 | 1,000,000 to 4,000,000 | Range used (alt to Viboud 2015) [3] |
| Influenza (H3N2) | pandemic | 1969 | 2,000,000 | 1,000,000 to 4,000,000 | Global estimate range [4] |
| SARS-CoV-2 (COVID-19) | pandemic | 2023 | 27,000,000 | 18,000,000 to 30,000,000 | Cumulative excess deaths (global) through ~late 2023 [5] |
| Ebola virus | BSL-4 | 2015 | 11,325 | 11,325 to 11,325 | West Africa outbreak 2014-2016 [6] |
| Marburg virus | BSL-4 | 2005 | 250 | 227 to 329 | Angola 2005 outbreak; deaths revised over time [7] |
| Nipah virus | BSL-4 | 1999 | 105 | 105 to 105 | Malaysia/Singapore 1998-1999 [8] |
| Hendra virus | BSL-4 | 2011 | 1 | 1 to 1 | Cumulative human deaths (Australia; no large outbreaks) Token value (1): no sustained human-to-human transmission; spillover-only. [9] |
| Crimean-Congo hemorrhagic fever virus | BSL-4 | 2022 | 40 | 27 to 52 | Largest recent outbreak benchmark (Iraq 2022) [10] |
| Lassa virus | BSL-4 | 2018 | 150 | 119 to 170 | Nigeria 2018 surge (note: endemic background ≈5,000/yr uncertain) [11] |
| Junín virus (Argentine HF) | BSL-4 | 1964 | 113 | 100 to 200 | Largest recorded Argentine HF epidemic mid-1960s; deaths ~113 [12] |
| Machupo virus (Bolivian HF) | BSL-4 | 1964 | 113 | 100 to 200 | San Joaquín 1963-1964 epidemic; deaths ~113 (reports vary) [13] |
| Guanarito virus (Venezuelan HF) | BSL-4 | 1995 | 140 | 120 to 200 | Deaths here approximate cumulative across multi-year outbreaks (largest single-year cluster smaller) [14] |
| Sabiá virus | BSL-4 | 1994 | 3 | 2 to 4 | Very few documented human cases; no large outbreak [15] |

| Agent | Category | Year | Best estimate | Low to high | Interpretation and sources |
|---|---|---|---|---|---|
| Lujo virus | BSL-4 | 2008 | 4 | 4 to 4 | Single nosocomial cluster Zambia-South Africa, 2008 (5 cases, 4 deaths) [16] |
| Chapare virus | BSL-4 | 2019 | 4 | 3 to 4 | 2019 Bolivia outbreak ~9 cases, 4 deaths; earlier 2003 single fatal case [17] |
| Omsk hemorrhagic fever virus | BSL-4 (often) | 1948 | 1 | 1 to 1 | Largest outbreak deaths not cleanly published; fatalities generally low Token value (1): no sustained human-to-human transmission; spillover-only. [18] |
| Kyasanur Forest disease virus | BSL-4 (often) | 1984 | 1 | 1 to 1 | Largest documented yearly tolls in 1983-1984; deaths 110 and 139 Token value (1): no sustained human-to-human transmission; spillover-only. [19] |
| Tick-borne encephalitis virus - Far Eastern | BSL-4 (some labs) | - | 1 | 1 to 1 | Subtype of TBEV; outbreak death counts vary by country; largest single events generally tens of deaths, not hundreds [20] |
| Tick-borne encephalitis virus - European | BSL-4 (some labs) | - | 1 | 1 to 1 | Subtype/region category; no distinct single-agent outbreak totals [21] |
| Herpes B (Macacine herpesvirus 1) | BSL-4 (laboratory work) | 1997 | 1 | 1 to 1 | No outbreaks; cumulative fatalities across decades; plotted as reference Token value (1): no sustained human-to-human transmission; spillover-only. [22] |

A value of 1 is explicitly identified as a plotting placeholder for Hendra, Omsk, Kyasanur Forest disease and herpes B. It is not an observed outbreak death total. The tick-borne encephalitis rows also require the source notes when interpreting their values. Years are reproduced as supplied; an unspecified year is not imputed. HF = hemorrhagic fever.

#### Supplementary Table S4. Risk thresholds across sectors

Sector summaries and numerical anchors are reproduced from the source workbook. Annual individual risks, lifetime risks, societal-risk curves and per-event criteria are different quantities and are not treated as interchangeable standards.

| Sector or domain | Named framework | Metric | Numerical anchor | Normalization and limitations |
|---|---|---|---|---|
| Major hazards (process industries) | UK HSE (R2P2/COMAH) | Public IR, FN | IR ≈ $10^{-6}$ (acceptable), $10^{-4}$ (intolerable); ALARP between | IR → expected fatalities only as illustrative bound (P×IR); not for compliance |
| External safety | Netherlands Bevi | Public IR, FN | IR contour $10^{-6}$/yr; FN societal criteria | As above |
| Seveso III (EU) implementations | Member-state practice | Public IR, FN | Typically IR $10^{-6}$/yr, $10^{-4}$/yr; FN curves | Country-specific details vary |
| Road/Rail tunnels | EU practice, Channel Tunnel, Italy (Gu@larp) | Public IR, FN | IR ~$10^{-6}$ acceptable, ~$10^{-4}$ intolerable; FN criteria | Group-specific thresholds per user class possible |
| Dam safety | USBR/USACE/ANCOLD | Public IR, FN | IR ~$10^{-6}$ to $10^{-4}$/yr; FN points (e.g., P(N≥1000)) | National details vary; directional |
| Nuclear (public risk goals) | U.S. NRC Safety Goals | Public risk to average individual | Early fatality ≤5 × $10^{-7}$/yr; latent cancer ≤2 × $10^{-6}$/yr | Accident-consequence framing |
| Space launch (range safety) | U.S. Range Safety | Expected casualty per event | E_c ≤ $10^{-6}$ per launch | E[fatalities/year] = launches/year × expected casualties/launch × fatal fraction |
| Aviation (ATM, TLS) | ICAO / EUR | System frequency | ~$10^{-9}$ to $10^{-8}$ fatal accidents per flight-hr | Convert to per aircraft-yr via (p × H × F) (assumptions explicit) |
| Aviation (system reliability) | Transport category aircraft | System frequency | Catastrophic system failure ~$10^{-9}$ per flight-hr | Not a public-risk limit; shows barrier reliability |
| Maritime | IMO FSA | Societal risk (FN) | Negligible ~$10^{-6}$; intolerable ~$10^{-3}$ (context-dependent) | Implemented per vessel/operation |
| Environmental (air toxics) | U.S. EPA residual risk | Lifetime individual risk | ~$10^{-6}$ acceptable; ~$10^{-4}$ presumptively unacceptable | Chronic, not accident; no annualization |
| Drinking water | EPA SDWA (MCL/MCLG) | Lifetime individual risk | Often targets ~$10^{-6}$ to $10^{-4}$ | Feasibility-bounded |
| Occupational (contrast) | OSHA jurisprudence | Worker lifetime risk | ~$10^{-3}$ lifetime significant (≈2.2 × $10^{-5}$/yr) | Shows ~$10^2$-$10^3$× public/worker gap |

IR = individual risk; FN = societal-risk frequency-number curve; ALARP = as low as reasonably practicable; ATM = air traffic management; TLS = target level of safety; FSA = formal safety assessment. $E_c$ denotes expected casualties. An illustrative conversion to fatalities requires the exposed population or activity frequency and the fatality fraction, as applicable.

Source documentation: the workbook identifies institutions and frameworks but does not give instrument editions, clause numbers or row-level source links for this table. The numerical anchors therefore retain the status of supplied illustrative summaries, rather than verified statements of current regulatory requirements.

### Supplementary Table S5. Subcritical pathogen calculations

#### Panel A. Input ranges and behavioral-response factors

The release rate $p_{init}$ is 0.0025 per year in every row. Low-high pairs preserve the supplied endpoints. $b_{max}$ = 0.9 and τ = 0.3.

| Agent | IFR: low to high | $R_{eff}$ before adjustment | Covert fraction: low to high | b at high IFR | b at low IFR |
|---|---|---|---|---|---|
| Ebola | 0.5 to 0.7 | 0.1 to 0.5 | 0.05 to 0.2 | 0.9 | 0.9 |
| Nipah | 0.4 to 0.75 | 0.4 to 0.9 | 0.1 to 0.45 | 0.9 | 0.9 |
| Lassa | 0.005 to 0.02 | 0.2 to 0.6 | 0.15 to 0.35 | 0.06 | 0.015 |
| CCHF | 0.01 to 0.3 | 0.05 to 0.3 | 0.05 to 0.2 | 0.9 | 0.03 |
| Marburg | 0.25 to 0.7 | 0.1 to 0.4 | 0.05 to 0.2 | 0.9 | 0.75 |
| Junin | 0.02 to 0.2 | 0.05 to 0.2 | 0.1 to 0.25 | 0.6 | 0.06 |
| Machupo | 0.15 to 0.3 | 0.05 to 0.2 | 0.1 to 0.25 | 0.9 | 0.45 |
| Guanarito | 0.2 to 0.3 | 0.02 to 0.1 | 0.1 to 0.25 | 0.9 | 0.6 |
| Chapare | 0.2 to 0.6 | 0.05 to 0.3 | 0.1 to 0.25 | 0.9 | 0.6 |
| Lujo | 0.4 to 0.8 | 0.1 to 0.4 | 0.1 to 0.25 | 0.9 | 0.9 |
| Sabiá | 0.15 to 0.5 | 0.02 to 0.1 | 0.1 to 0.25 | 0.9 | 0.45 |
| MERS | 0.1 to 0.35 | 0.1 to 0.4 | 0.05 to 0.2 | 0.9 | 0.3 |

IFR = infection fatality ratio, expressed as a fraction. The behavioral-response factors are dimensionless. The workbook labels the pre-adjustment transmission columns R_eff_low and R_eff_high; those labels are retained here.

Behavioral adjustment: $b(IFR) = b_{max} \min(1, IFR/\tau)$; $R_{eff,adj} = R_{eff}[c + (1 - c)(1 - b)]$, where c is the covert fraction. The low and high scenario pairings are those used in the workbook.

#### Supplementary Table S5 (continued)

##### Panel B. Adjusted transmission, expected harm and assigned level

| Agent | Adjusted $R_{eff}$: low to high | Expected fatalities/year: low to high | SSL | Outbreak size (supplied) |
|---|---|---|---|---|
| Ebola | 0.0145 to 0.14 | $1.3 \times 10^{-3}$ to $2.0 \times 10^{-3}$ | 3 | 11,325 |
| Nipah | 0.076 to 0.4545 | $1.1 \times 10^{-3}$ to $3.4 \times 10^{-3}$ | 3 | 105 |
| Lassa | 0.1898 to 0.59415 | $1.5 \times 10^{-5}$ to $1.2 \times 10^{-4}$ | 2 | 150 |
| CCHF | 0.00725 to 0.2928 | $2.5 \times 10^{-5}$ to $1.1 \times 10^{-3}$ | 3 | 40 |
| Marburg | 0.0145 to 0.16 | $6.3 \times 10^{-4}$ to $2.1 \times 10^{-3}$ | 3 | 250 |
| Junin | 0.023 to 0.191 | $5.1 \times 10^{-5}$ to $6.2 \times 10^{-4}$ | 2 | 113 |
| Machupo | 0.0095 to 0.1325 | $3.8 \times 10^{-4}$ to $8.6 \times 10^{-4}$ | 2 | 113 |
| Guanarito | 0.0038 to 0.055 | $5.0 \times 10^{-4}$ to $7.9 \times 10^{-4}$ | 2 | 140 |
| Chapare | 0.0095 to 0.165 | $5.0 \times 10^{-4}$ to $1.8 \times 10^{-3}$ | 3 | 4 |
| Lujo | 0.019 to 0.13 | $1.0 \times 10^{-3}$ to $2.3 \times 10^{-3}$ | 3 | 4 |
| Sabiá | 0.0038 to 0.06625 | $3.8 \times 10^{-4}$ to $1.3 \times 10^{-3}$ | 3 | 3 |
| MERS | 0.0145 to 0.304 | $2.5 \times 10^{-4}$ to $1.3 \times 10^{-3}$ | 3 | 93 |

Expected harm: $E[harm] = p_{init} \times IFR / (1 - R_{eff,adj})$. Expected-harm values are displayed to two significant figures; the underlying spreadsheet values are not altered. SSL assignments are reproduced as supplied. The "Outbreak size" field is retained under its original label because the workbook does not define its unit consistently in the column header.

Calculation provenance: the supplied worksheet combines formulas and stored numerical outputs. This PDF reports its saved values. It does not imply that every output recalculates when inputs change.

#### Supplementary Table S6. Evidence and classification rationale

Agent classifications, stated lower bounds and decisions are reproduced from the workbook, including the Hendra and SARS-CoV-1 exceptions. Sources from the two original source columns are consolidated into numbered references.

| Agent | Classification basis | Decision | M (lower bound) | Rationale and sources |
|---|---|---|---|---|
| Nipah | Respiratory-capable, spillover count | Subcritical → finite-chain | ≥183 | Bangladesh documents 183 independent spillovers (2001-2018); no sustained global transmission therefore rule-of-3 bound puts p(pandemic in high-income settings) ≤ 1.64%. [23] |
| MERS-CoV | Respiratory-capable, spillover count | Subcritical → finite-chain | ≥60 (conservative) | Since 2012, ~2,600+ confirmed cases; transmission largely nosocomial; community (R<1) and non-sustained spread therefore many independent introductions (M much greater than 60). [24] |
| Ebola (all spp.) | Non-respiratory | Subcritical → finite-chain | ≥30 | Body-fluid/close-contact transmission; many outbreaks, but efficient casual-contact spread in HICs not seen. [25] |
| Marburg | Non-respiratory | Subcritical → finite-chain | ≥18 | Dozens of independent episodes since 1967; human spread is close-contact/nosocomial; no sustained community chains in HICs. [26] |
| Lassa | Non-respiratory | Subcritical → finite-chain | - | Rodent-borne; person-to-person mostly nosocomial; casual contact not a driver in HICs. (Endemic-M not comparable.) [27] |
| CCHF | Non-respiratory | Subcritical → finite-chain | much greater than 100 (global) | Extensive global experience; H2H largely nosocomial (review tallies 158 nosocomial cases, 1953-2016); community propagation not sustained. [28] |
| Junín (AHF) | Non-respiratory | Subcritical → finite-chain | - | Rodent-borne; H2H very rare, nosocomial possible; household SAR ≈ zero in reports. [29] |
| Guanarito (Venezuelan HF) | Non-respiratory | Subcritical → finite-chain | - | Endemic zoonosis; no widespread H2H; nosocomial described; community chains not sustained. [30] |
| Herpes B (Macacine herpesvirus 1) | Non-H2H | Subcritical → finite-chain | - | ~50 documented human infections since 1932; 1 confirmed H2H (spousal); remaining cases macaque → human therefore community (R ≈ 0). [31] |
| Kyasanur Forest disease (KFD) | Non-H2H, vector-borne | Subcritical → finite-chain | - | Tick-borne; humans are dead-end hosts; no H2H despite large seasonal burdens. [32] |
| Omsk hemorrhagic fever (OHF) | Non-H2H, vector/rodent | Subcritical → finite-chain | - | Tick/rodent-associated; no human-to-human transmission reported across decades. [33] |

| Agent | Classification basis | Decision | M (lower bound) | Rationale and sources |
|---|---|---|---|---|
| Tick-borne encephalitis virus (TBEV) | Non-H2H, tick/food-borne | Subcritical → finite-chain | - | Tick/food-borne; not transmitted person-to-person (rare special routes). [34] |
| Machupo (BHF) | Non-respiratory | Subcritical → finite-chain | ~5-6 | Evidence too sparse to certify non-resp with audit thresholds; nosocomial H2H reported but not sustaining epidemics. [35] |
| Sabiá | Non-respiratory | Subcritical → finite-chain | <5 | Too few human cases to certify non-resp under thresholds. [36] |
| Chapare | Non-respiratory | Subcritical → finite-chain | 2-3 | Small HCW cluster; H2H shown but evidence base too small to certify non-resp. [37] |
| Lujo | Non-respiratory | Subcritical → finite-chain | 1 | Single nosocomial cluster; too sparse for certification. [38] |
| Hendra | Few spillovers | SSL-5 (default) | - | Henipavirus clade (resp-capable); no H2H documented, but human data are sparse-treat as high until evidence accrues. [39] |
| SARS-CoV-1 | Demonstrated R0 > 1 | SSL-5 (default) | - | Supercritical without controls (pre-control (R0 ~ 2-3), strong superspreading). [40] |

M denotes the reported lower bound on independent introductions or episodes; a dash indicates no comparable bound was provided. HIC = high-income country; H2H = human-to-human; HCW = health-care worker; SAR = secondary attack rate; AHF/BHF = Argentine/Bolivian hemorrhagic fever. The source table mixes different evidence types; its M values are not all directly comparable.

### Supplementary Table S7. Supercritical and uncertain scenarios

#### Panel A. Parameters and expected harm

$p_{init}$ = 0.0025 per year for every scenario. Entries reproduce the workbook's two-significant-figure presentation table; low-high pairs are the supplied endpoints.

| Scenario | $\pi_{super}$ | R: low to high | k: low to high | $P_{amp}$: low to high | C: low to high | E[harm]/year: low to high |
|---|---|---|---|---|---|---|
| 1918-class influenza (today) | 0.1 | 2 to 2.4 | 1 to 2 | 0.5 to 0.71 | 20,000,000 to 80,000,000 | $2.5 \times 10^3$ to $1.4 \times 10^4$ |
| 1957 H2N2 | 0.1 | 1.5 to 2 | 1 to 2 | 0.33 to 0.62 | 1,500,000 to 6,000,000 | $1.2 \times 10^2$ to $9.3 \times 10^2$ |
| 1968 H3N2 | 0.01 | 1.4 to 1.9 | 1 to 2 | 0.29 to 0.59 | 1,500,000 to 6,000,000 | 11 to 88 |
| 1977 H1N1 | 0.1 | 1.4 to 1.9 | 1 to 2 | 0.29 to 0.59 | 200,000 to 700,000 | 14 to $1 \times 10^2$ |
| SARS-CoV-1 | 0.1 | 2 to 3 | 0.1 to 0.3 | 0.11 to 0.37 | 4,000,000 to 20,000,000 | $1.1 \times 10^2$ to $1.9 \times 10^3$ |
| Antibody-escaping SARS-CoV-2 | 1 | 6 to 8 | 0.1 to 0.2 | 0.24 to 0.44 | 1,500,000 to 6,000,000 | $9 \times 10^2$ to $6.7 \times 10^3$ |
| Nipah - enhanced (R=1.6) | 1 | 1.4 to 1.8 | 0.1 to 0.3 | 0.057 to 0.23 | 10,000,000 to 40,000,000 | $1.4 \times 10^3$ to $2.3 \times 10^4$ |
| Nipah - mutant to R=1.6 | 0.001 | 1.4 to 1.8 | 0.1 to 0.3 | 0.057 to 0.23 | 10,000,000 to 40,000,000 | 1.4 to 23 |
| Human-infecting bat CoV spreads | 0.005 | 2.5 to 3.5 | 0.1 to 0.2 | 0.14 to 0.31 | 10,000,000 to 40,000,000 | 17 to $1.6 \times 10^2$ |
| Variola major | 0.005 | 3.5 to 6 | 0.5 to 1 | 0.54 to 0.83 | 30,000,000 to 600,000,000 | $2 \times 10^2$ to $6.3 \times 10^3$ |

$\pi_{super}$ is the assigned probability for the stated scenario; R is the reproduction number; k is the dispersion parameter; $P_{amp}$ is amplification probability; C is fatalities conditional on amplification. E[harm] = $p_{init} \times \pi_{super} \times P_{amp} \times C$. Display rounding can prevent exact reproduction by multiplying the printed inputs.

These ranges are conditional scenario ranges, not statistical confidence intervals. They retain the workbook assumptions and do not resolve differences between the workbook and the manuscript figures.

### Supplementary Table S7 (continued)

#### Panel B. Scenario justifications

| Scenario | $\pi_{super}$ | Supplied justification |
|---|---|---|
| 1918-class influenza (today) | 0.1 | 10% prior that population immunity is insufficient to prevent global spread |
| 1957 H2N2 | 0.1 | 10% prior that population immunity is insufficient to prevent global spread |
| 1968 H3N2 | 0.01 | 1% prior that population immunity is insufficient to prevent global spread |
| 1977 H1N1 | 0.1 | 10% prior that population immunity is insufficient to prevent global spread |
| SARS-CoV-1 | 0.1 | Containment failure before export to LIC megacity slums |
| Antibody-escaping SARS-CoV-2 | 1 | Escape variant is created to test viral immune evasion model |
| Nipah - enhanced (R=1.6) | 1 | Assumes a lab performs an enhanced-transmission study on Nipah |
| Nipah - mutant to R=1.6 | 0.001 | 0.1% chance an experimental mutation achieves equivalent enhancement |
| Human-infecting bat CoV spreads | 0.005 | 1,200 to 6,000 bat CoVs; ~10% infect humans; 2 known supercritical examples: 2/400 |
| Variola major | 0.005 | Escape into LIC megacity slums before control |

LIC = low-income country; CoV = coronavirus. The numerical values and scenario justifications above are retained from the workbook. Numerical iteration details are retained in its Supercritical-raw worksheet.

### Calculation methods and workbook

#### Subcritical calculations

The adjusted transmission and expected-harm equations are shown with Table S5. The low endpoint combines low IFR, low pre-adjustment transmission, low covert fraction and the behavioral response evaluated at high IFR. The high endpoint uses the opposite supplied pairing. This describes the worksheet convention; it is not a statistical interval calculation.

#### Supercritical calculations

The manuscript models early transmission using a negative-binomial branching process. With $p = k/(k + R)$, the extinction probability q satisfies $q = [p / (1 - (1 - p)q)]^k$, and $P_{amp} = 1 - q$. The relevant solution is the extinction probability in [0, 1], rather than the trivial $q = 1$ root when $R > 1$.

The workbook records $q_0 = 0.99$, 25 displayed iteration steps, and separate numerical endpoints for the low and high parameter sets. The Methods state a tolerance of $10^{-8}$. The saved iteration grids and endpoint values alone do not demonstrate that tolerance for every row; they are reproduced as stored calculations, not independently re-run here.

#### Reproducibility and numerical precision

Tables S5 and S7 in this PDF report the values saved in the supplied workbook. Some downstream values and numerical iterations are constants rather than formulas. The workbook should be consulted for unrounded values, parameter provenance and intermediate results; changing inputs will not necessarily refresh every stored output.

Supplementary Information and the electronic workbook are available from Zenodo:

https://doi.org/10.5281/zenodo.23745003

#### Supplementary source references

Numbering applies to Tables S3 and S6. Source descriptions are retained from the workbook; URLs are clickable.

**[1]** Influenza (H1N1). Our World in Data - Historical Pandemics
https://ourworldindata.org/historical-pandemics

**[2]** Influenza (H1N1). Rozo et al 10.1128/mBio.01013-15
https://doi.org/10.1128/mBio.01013-15

**[3]** Influenza (H2N2). Our World in Data - Historical Pandemics
https://ourworldindata.org/historical-pandemics

**[4]** Influenza (H3N2). Our World in Data - Historical Pandemics
https://ourworldindata.org/historical-pandemics

**[5]** SARS-CoV-2 (COVID-19). Our World in Data - Excess mortality
https://ourworldindata.org/excess-mortality-covid
https://ourworldindata.org/grapher/cumulative-excess-deaths-covid

**[6]** Ebola virus. CDC MMWR overview of 2014-16 West Africa epidemic; CDC Ebola Outbreak History page
https://www.cdc.gov/mmwr/volumes/65/su/su6503a2.htm
https://www.cdc.gov/ebola/outbreaks/index.html

**[7]** Marburg virus. WHO DON (Apr 2005); CIDRAP (Nov 2005 revision to 227 deaths)
https://www.who.int/emergencies/disease-outbreak-news/item/2005_04_22-en
https://www.cidrap.umn.edu/ebola/angola-declares-worst-marburg-outbreak-over

**[8]** Nipah virus. Looi 2007 / PubMed; Ang et al. 2018 / PMC review
https://pubmed.ncbi.nlm.nih.gov/19108397/
https://pmc.ncbi.nlm.nih.gov/articles/PMC5971524/

**[9]** Hendra virus. WHO Hendra factsheet
https://www.who.int/health-topics/hendra-virus-infection

**[10]** Crimean-Congo hemorrhagic fever virus. WHO DON on Iraq 2022; Alhilfi 2023 (largest outbreak in Iraq)
https://www.who.int/emergencies/disease-outbreak-news/item/2022-DON386
https://pmc.ncbi.nlm.nih.gov/articles/PMC9931897/

**[11]** Lassa virus. WHO DON (Mar 2018 Nigeria); NCDC SitRep #52 (Dec 31, 2018)
https://www.who.int/emergencies/disease-outbreak-news/item/23-march-2018-lassa-fever-nigeria-en
https://reliefweb.int/report/nigeria/2018-lassa-fever-outbreak-nigeria-ncdc-situation-report-52-31-december-2018

**[12]** Junín virus (Argentine HF). Machupo/Junín outbreak summaries (SciDirect; review)
https://www.sciencedirect.com/topics/agricultural-and-biological-sciences/junin-virus
https://www.sciencedirect.com/topics/medicine-and-dentistry/bolivian-hemorrhagic-fever

**[13]** Machupo virus (Bolivian HF). PHAC Pathogen Safety Data Sheet (2012); Review of BHF outbreaks (SciDirect)
https://www.canada.ca/en/public-health/services/laboratory-biosafety-biosecurity/pathogen-safety-data-sheets-risk-assessment/machupo-virus.html
https://www.sciencedirect.com/topics/medicine-and-dentistry/bolivian-hemorrhagic-fever

**[14]** Guanarito virus (Venezuelan HF). Silva-Ramos 2022 review (CFR ~33%)
https://pmc.ncbi.nlm.nih.gov/articles/PMC9110938/

**[15]** Sabiá virus. CDC/WHO/Reviews; sporadic cases only; CDC weekly notifiable disease tables (surveillance)
https://ndc.services.cdc.gov/case-definitions/new-world-arenavirus-sabia-virus/
https://stacks.cdc.gov/view/cdc/232133

**[16]** Lujo virus. CDC Lujo virus page; GIDEON summary
https://www.cdc.gov/lujo-fever/about/index.html
https://www.gideononline.com/blogs/lujo-hemorrhagic-fever-luhf/

**[17]** Chapare virus. WHO DON (Jan 2025); CDC Chapare page (archived mirror)
https://www.who.int/emergencies/disease-outbreak-news/item/2025-DON553
https://www.restoredcdc.org/www.cdc.gov/chapare/about/index.html

**[18]** Omsk hemorrhagic fever virus. CDC OHF overview; CFR 0.5-3%
https://www.cdc.gov/omsk-fever/about/index.html
https://stacks.cdc.gov/view/cdc/24421/cdc_24421_DS1.pdf

**[19]** Kyasanur Forest disease virus. Karnataka DHFWS Operational Manual (reports 110 deaths in 1983 and 139 in 1984); Peer-reviewed overviews of KFD outbreaks
https://monkeyfeverrisk.ceh.ac.uk/sites/default/files/2022-05/Operational-Manual-Kyasanur-Forest-Disease-DHFWS-2020-compressed.pdf
https://pmc.ncbi.nlm.nih.gov/articles/PMC7007137/

**[20]** Tick-borne encephalitis virus - Far Eastern. ECDC/Reviews on TBE CFR (up to ~35% for Far Eastern)
https://www.ecdc.europa.eu/en/tick-borne-encephalitis/facts/factsheet
https://www.ecdc.europa.eu/en/tick-borne-encephalitis

**[21]** Tick-borne encephalitis virus - European. ECDC factsheet (mortality ~0.5-2%); EU 2012-2020: 93 deaths total (Van Heuverswyn 2023)
https://www.ecdc.europa.eu/en/tick-borne-encephalitis/facts/factsheet
https://pmc.ncbi.nlm.nih.gov/articles/PMC10021474/

**[22]** Herpes B (Macacine herpesvirus 1). CDC Clinical Overview (50 human cases, 21 deaths); MMWR 1997 lab exposure fatality; reviews
https://www.cdc.gov/herpes-b-virus/hcp/clinical-overview/index.html
https://www.cdc.gov/mmwr/preview/mmwrhtml/00056008.htm

**[23]** Nipah. Bangladesh spillover compilation (183 events, 2001-2018); CDC overview; WHO factsheet
https://www.ncbi.nlm.nih.gov/pmc/articles/PMC5971524/
https://www.cdc.gov/vhf/nipah/
https://www.who.int/health-topics/nipah-virus

**[24]** MERS-CoV. WHO MERS fact sheet/overview; CDC MERS; Review of transmission (community (R<1), nosocomial clusters); Korea 2015 nosocomial outbreak summary
https://www.who.int/health-topics/mers
https://www.cdc.gov/coronavirus/mers/about/index.html
https://www.ncbi.nlm.nih.gov/pmc/articles/PMC7391875/
https://www.ncbi.nlm.nih.gov/pmc/articles/PMC4543038/

**[25]** Ebola (all spp.). CDC outbreak chronology; CDC At-a-Glance
https://www.cdc.gov/vhf/ebola/history/chronology.html
https://www.cdc.gov/vhf/ebola/pdf/ebola-factsheet.pdf

**[26]** Marburg. WHO Marburg fact sheet; CDC "History of Marburg Outbreaks"
https://www.who.int/news-room/fact-sheets/detail/marburg-virus-disease
https://www.cdc.gov/vhf/marburg/outbreaks/history.html

**[27]** Lassa. WHO Lassa factsheet; CDC Lassa
https://www.who.int/news-room/fact-sheets/detail/lassa-fever
https://www.cdc.gov/vhf/lassa/

**[28]** CCHF. Nosocomial review (158 cases); WHO CCHF factsheet
https://pubmed.ncbi.nlm.nih.gov/31992842/
https://www.who.int/health-topics/crimean-congo-haemorrhagic-fever

**[29]** Junín (AHF). UK HCID (Junín); Review (Viruses)
https://www.gov.uk/guidance/high-consequence-infectious-diseases-hcid#arenaviridae
https://www.ncbi.nlm.nih.gov/pmc/articles/PMC8706732/

**[30]** Guanarito (Venezuelan HF). 2022 review (GTOV); BMC Microbiology 2024 review
https://www.ncbi.nlm.nih.gov/pmc/articles/PMC9110938/
https://bmcmicrobiol.biomedcentral.com/articles/10.1186/s12866-024-03128-3

**[31]** Herpes B (Macacine herpesvirus 1). CDC Clinical Overview; MMWR 1997 lab exposure
https://www.cdc.gov/herpes-b-virus/hcp/clinical-overview/index.html
https://www.cdc.gov/mmwr/preview/mmwrhtml/00056008.htm

**[32]** Kyasanur Forest disease (KFD). Operational manual (India); Review
https://monkeyfeverrisk.ceh.ac.uk/sites/default/files/2022-05/Operational-Manual-Kyasanur-Forest-Disease-DHFWS-2020-compressed.pdf
https://www.ncbi.nlm.nih.gov/pmc/articles/PMC7007137/

**[33]** Omsk hemorrhagic fever (OHF). CDC OHF; Historical review
https://www.cdc.gov/omsk-fever/about/index.html
https://stacks.cdc.gov/view/cdc/24421/cdc_24421_DS1.pdf

**[34]** Tick-borne encephalitis virus (TBEV). CDC TBE; ECDC factsheet
https://www.cdc.gov/tick-borne-encephalitis/index.html
https://www.ecdc.europa.eu/en/tick-borne-encephalitis/facts/factsheet

**[35]** Machupo (BHF). Review/PSDS; Review
https://www.canada.ca/en/public-health/services/laboratory-biosafety-biosecurity/pathogen-safety-data-sheets-risk-assessment/machupo-virus.html
https://www.sciencedirect.com/topics/medicine-and-dentistry/bolivian-hemorrhagic-fever

**[36]** Sabiá. CDC case definition/overview; Review
https://ndc.services.cdc.gov/case-definitions/new-world-arenavirus-sabia-virus/
https://stacks.cdc.gov/view/cdc/232133

**[37]** Chapare. NEJM 2020/2022 Chapare reports; WHO DON 2025
https://www.nejm.org/doi/full/10.1056/NEJMoa2004194
https://www.who.int/emergencies/disease-outbreak-news/item/2025-DON553

**[38]** Lujo. CDC Lujo; GIDEON summary
https://www.cdc.gov/lujo-fever/about/index.html
https://www.gideononline.com/blogs/lujo-hemorrhagic-fever-luhf/

**[39]** Hendra. WHO Hendra; NSW Health fact sheet
https://www.who.int/health-topics/hendra-virus-infection
https://www.health.nsw.gov.au/Infectious/factsheets/Pages/hendra_virus.aspx

**[40]** SARS-CoV-1. Lipsitch et al., Science 2003; Lloyd-Smith et al., Nature 2005
https://www.science.org/doi/10.1126/science.1086616
https://www.nature.com/articles/nature04153
